# Didemnin B and ternatin-4 inhibit conformational changes in eEF1A required for aminoacyl-tRNA accommodation into mammalian ribosomes

**DOI:** 10.1101/2022.08.01.502281

**Authors:** Manuel F. Juette, Jordan D. Carelli, Emily J. Rundlet, Alan Brown, Sichen Shao, Angelica Ferguson, Michael R. Wasserman, Mikael Holm, Jack Taunton, Scott C. Blanchard

## Abstract

Rapid and accurate mRNA translation requires efficient codon-dependent delivery of the correct aminoacyl-tRNA (aa-tRNA) to the ribosomal A site. In mammals, this fidelity-determining reaction is facilitated by the GTPase elongation factor-1 alpha (eEF1A), which escorts aa-tRNA as an eEF1A(GTP)-aa-tRNA ternary complex into the ribosome. Two structurally unrelated cyclic peptides didemnin B and ternatin-4 bind to the eEF1A(GTP)-aa-tRNA ternary complex and inhibit translation. Here, we employ single-molecule fluorescence imaging and cryogenic electron microscopy to determine how these natural products inhibit translational elongation on mammalian ribosomes. By binding to a common allosteric site on eEF1A, didemnin B and ternatin-4 trap eEF1A in its GTPase-activated conformation, preventing aa-tRNA accommodation on the ribosome. We also show that didemnin B and ternatin-4 exhibit distinct effects on aa-tRNA selection that inform on observed disparities in their inhibition efficacies and physiological impacts. These integrated findings highlight the potential of single-molecule methods to reveal how distinct natural products differentially impact the human translation mechanism.

## Introduction

Translation of the genetic code from mRNA into protein is a multi-step process catalyzed by the two-subunit ribosome (80S in eukaryotes) in coordination with translational GTPases (Behrmann et al., 2015). Each translation step is regulated by signaling pathways linked to cell growth, differentiation, nutrient sensing, and homeostatic quality control. Protein synthesis status is thus a central hub for sensing cellular stress. Dysregulated protein synthesis plays a role in several human diseases and is a therapeutic vulnerability in cancer and viral infection (Bhat et al., 2015; Hoang et al., 2021; Xu and Ruggero, 2020).

The elongation cycle in eukaryotic protein synthesis begins with the binding of a ternary complex of the highly conserved, three-domain (DI-III) eukaryotic elongation factor-1 alpha (eEF1A), GTP, and aminoacyl-tRNA (aa-tRNA) to the Aminoacyl (A) site at the leading edge of the 80S ribosome (Abbas et al., 2015). Base-pairing interactions within the small subunit (SSU) between the A-site mRNA codon and a cognate aa-tRNA anticodon trigger a sequence of structural rearrangements that dock eEF1A at the large subunit (LSU) GTPase activating center (GAC). There, the GAC triggers eEF1A to hydrolyze GTP, ultimately driving eEF1A dissociation and accommodation of the aa-tRNA 3’-CCA end into the LSU peptidyl transferase center (PTC) (Budkevich et al., 2014; Ferguson et al., 2015; Voorhees et al., 2010). Once fully accommodated, aa-tRNA undergoes a peptide-bond forming, condensation reaction that extends the nascent polypeptide by one amino acid, generating a pre-translocation ribosome complex. Ensuing conformational processes within the ribosome enable engagement by eEF2, which catalyzes mRNA and tRNA translocation to complete the elongation cycle (Noller et al., 2017). Processive elongation reactions repeat over hundreds to thousands of mRNA codons to synthesize proteins.

Multiple natural products identified in phenotypic screens for anticancer or other biological activities directly target eEF1A (Carelli et al., 2015; Crews et al., 1994; Klein et al., 2021; Krastel et al., 2015; Lindqvist et al., 2010; Sun et al., 2021). Of these, didemnin B (henceforth “didemnin”) and its variants have been studied most extensively, including clinical trials for the treatment of specific cancer indications (Kucuk et al., 2000; Mittelman et al., 1999; Taylor et al., 1998; Vera and Joullié, 2002; Williamson et al., 1995) and severe acute respiratory syndrome coronavirus 2 (SARS-CoV-2 or COVID-19) (White et al., 2021; Yan et al., 2021). A cryogenic microscopy (cryo-EM) reconstruction of elongating rabbit reticulocyte lysate ribosomes revealed that didemnin traps eEF1A on the ribosome during aa-tRNA selection immediately after GTP hydrolysis and inorganic phosphate (P_i_) release, binding between DI and DIII of eEF1A (Shao et al., 2016). Ternatin-4, a cyclic peptide chemically unrelated to didemnin (**Figure 1––figure supplement 1**), also targets eEF1A competitively with didemnin to inhibit translation elongation (Carelli et al., 2015). A homozygous DIII point mutation (A399V) in eEF1A adjacent to the didemnin binding site confers nearly complete protection against the antiproliferative effects of ternatin-4 in HCT116 cells (Carelli et al., 2015). By contrast, cells harboring mutant eEF1A(A399V) are only partially resistant to didemnin (Krastel et al., 2015).

Here, we use single-molecule fluorescence resonance energy transfer (smFRET) imaging and comparative cryo-EM structural analysis of partially and fully reconstituted mammalian ribosome complexes to elucidate the effects of didemnin and ternatin-4 on aa-tRNA selection. We show that despite sharing the same allosteric binding site on eEF1A, didemnin and ternatin-4 differentially perturb the conformational dynamics of ribosome-associated eEF1A(GTP)-aa-tRNA ternary complex in ways that correlate with their effects on cellular growth and protein synthesis. These observations shed light on how these drugs impact the rate-determining conformational changes in eEF1A that govern aa-tRNA accommodation prior to peptide-bond formation.

## Results

### Didemnin and ternatin-4 inhibit aa-tRNA accommodation

We set out to examine the mechanistic impacts of didemnin and ternatin-4 on the process of aa-tRNA selection on human ribosomes using a smFRET platform that enables interrogation of purified, reconstituted human translation elongation reactions (Ferguson et al., 2015). This platform has been successfully employed to dissect kinetic and structural features of the eukaryotic elongation cycle as well as its modulation by plant and microbial natural products (Flis et al., 2018; McMahon et al., 2019; Pellegrino et al., 2019; Prokhorova et al., 2017).

Analogous to prior investigations of the bacterial aa-tRNA selection mechanism (Blanchard et al., 2004a; Geggier et al., 2010; Juette et al., 2016), this smFRET platform monitors the change in distance between fluorescently labeled incoming A-site aa-tRNA and P-site tRNA. These measurements yield quantitative structural and kinetic data that define the aa-tRNA selection mechanism, including the rates of eEF1A(GTP)-aa-tRNA ternary complex binding and the stepwise progression of aa-tRNA through distinct codon-recognition (CR), GTPase-activated (GA), and fully accommodated (AC) positions within the A site of the ribosome *en route* to peptide bond formation (**Figure 1A**) (Ferguson et al., 2015; Geggier et al., 2010).

**Figure 1.**
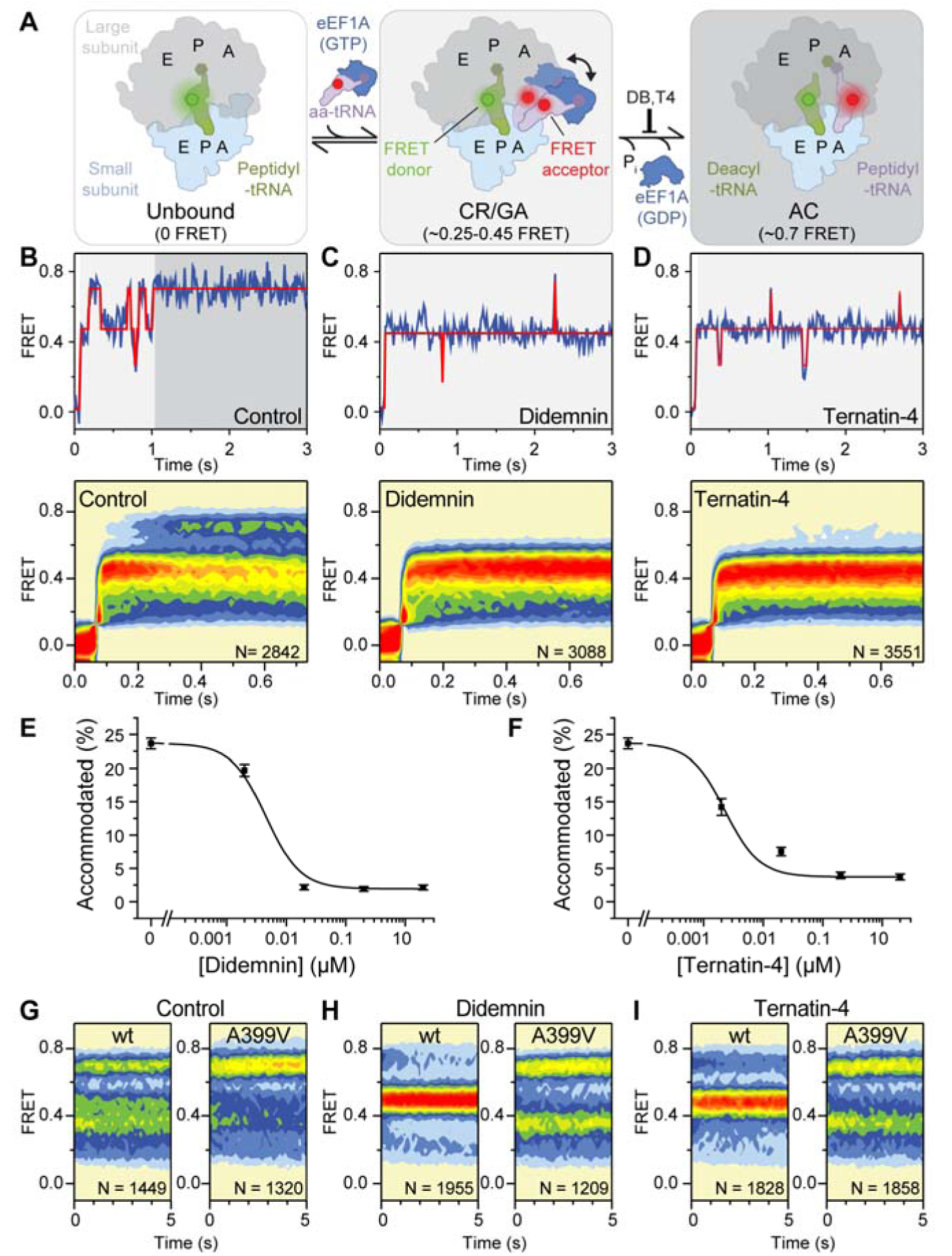
Mechanism of didemnin and terrnatin-4 inhibition revealed by smFRET. (**A**) Schematic of the experimental setup. Acceptor Cy5-labeled eEF1A(GTP)-aa-tRNA ternary complex is delivered to 80S initiation complexes with donor Cy3-labeled P-site tRNA (left). Codon recognition (CR; low FRET) leads to a pre-accommodated GTPase-activated (GA), mid-FRET state (center). Accommodation (AC) and peptide-bond formation produce a pre-translocation complex (right), which samples classical (high-FRET) and hybrid (mid-FRET) conformations in equilibrium. (**B**-**D**, *top*) Representative smFRET traces and (*bottom*) post-synchronized population histograms of N traces of accommodation dynamics of pre-steady state reactions in the presence of (B) DMSO (control) or 20 µM (C) didemnin or (D) ternatin-4. Shading behind traces and histograms indicates the predominant state assignment as described in (A; white, unbound; light gray, CR/GA; dark gray, accommodated). (**E**, **F**) Dose-response curves of the accommodated fraction in the presence of (E) didemnin or (F) ternatin-4. Error bars: s.e.m. from 1000 bootstrap samples. (**G**-**I**) Population histograms of N traces for steady-state reactions, formed with ternary complex containing recombinant eEF1A (WT or A399V). See also Figure 1—figure supplement 1-3.

Functional human 80S initiation complexes (ICs) were reconstituted from ribosomal subunits isolated from HEK293T cells, synthetic mRNA, and fluorescently labeled initiator tRNA (Ferguson et al., 2015) (Methods). Human ICs were surface-tethered within passivated microfluidic flow cells by a biotin-streptavidin bridge on the 5’ end of the mRNA (Juette et al., 2016). Ternary complex was formed with eEF1A purified from rabbit reticulocyte lysate (identical in primary sequence to human eEF1A1), fluorescently labeled Phe-tRNA^Phe^, and GTP (Methods). Pre-formed ternary complex was stopped-flow delivered to the immobilized ICs while imaging in real time to assess the specific effects of didemnin and ternatin-4 (Methods).

In the absence of inhibitor, the process of aa-tRNA selection was accompanied by a stepwise progression of aa-tRNA into the A site, which ultimately achieved a stable AC state, characterized by high (∼0.7) FRET efficiency (**Figure 1A**, **B**), as shown previously (Ferguson et al., 2015). As has been described by smFRET and structural studies of bacterial translation (Geggier et al., 2010; Munro et al., 2007; Rundlet et al., 2021; Whitford et al., 2010), the AC state is consistent with a classical (A/A) peptidyl-tRNA position within the pre-translocation complex site after peptide bond formation. Consistent with prior investigations in bacteria (Blanchard et al., 2004b; Geggier et al., 2010; Morse et al., 2020) and mammals (Ferguson et al., 2015), entrance into the final AC state at the end of aa-tRNA selection was accompanied by transient, reversible movements through two key intermediates in the aa-tRNA selection process characterized by low- (CR, ∼0.2) and intermediate- (GA, ∼0.45) FRET efficiencies (**Figure 1A**, **B**).

Quantitative investigations of bacterial translation have revealed that CR to GA and GA to AC transitions reflect conformational sampling of aa-tRNA between distinct positions within the A site that are directly related to the two-step kinetic proofreading mechanism underpinning decoding fidelity (Blanchard et al., 2004a; Geggier et al., 2010; Ieong et al., 2016; Morse et al., 2020; Whitford et al., 2010). The relatively rapid, initial selection phase of aa-tRNA selection prior to GTP hydrolysis is comprised of rapid reversible transitions between CR and GA-like states. The relatively slow, proofreading phase of aa-tRNA selection after GTP hydrolysis is comprised of reversible transitions between GA-like and AC-like states (Geggier et al., 2010; Morse et al., 2020; Whitford et al., 2010). Hence, processes following GTP hydrolysis are rate-limiting to the aa-tRNA selection mechanism in both bacteria and mammals.

In the presence of 20 µM didemnin, ribosome complexes efficiently stalled in a long-lived, GA-like (∼0.45 FRET efficiency) state (**Figure 1C**). These findings are consistent with the cryo-EM structure of didemnin-stalled elongation complexes isolated from rabbit reticulocyte lysate, in which peptidyl-tRNA was “classically” positioned within the P site (“P/P” configuration) and aa-tRNA adopted a bent “A/T” configuration bound to eEF1A at the subunit interface (Shao et al., 2016). This structure revealed that eEF1A was trapped in an active, GTP-bound conformation even after GTP hydrolysis and P_i_ release. These findings are also in line with extensive investigations of antibacterial compounds (i.e. kirromycin) that bind directly to EF-Tu, the bacterial homolog of eEF1A, to trap ternary complex on the bacterial ribosome in analogous GA-like states after GTP hydrolysis and P_i_ release (Fischer et al., 2015; Schmeing et al., 2009).

In the presence of 20 µM ternatin-4, we obtained results highly similar to those observed with didemnin (**Figure 1D**). Hence, both didemnin and ternatin-4 trap eEF1A on the leading edge of the human ribosome in intermediate states of aa-tRNA selection by slowing processes required for aa-tRNA accommodation from a timescale of hundreds of milliseconds to minutes. The finding that both structurally and chemically distinct molecules trap aa-tRNA in a GA-like state argues that they similarly inhibit the proofreading stage of aa-tRNA selection. More specifically, they likely slow the rate-limiting conformational changes within eEF1A(GTP/GDP)- aa-tRNA complex that allow aa-tRNA accommodation.

Quantifying the fraction of smFRET trajectories that rapidly reached the AC state, thus considered molecules that escaped drug inhibition, revealed similar dose-dependent inhibition profiles for didemnin and ternatin-4 (**Figure 1E**, **F**), with IC_50_ values of 4.5 ± 0.6 nM and 2.3 ± 0.4 nM, respectively. The dose-dependent effects of two ternatin variants showed that ternatin-2 was completely inactive and ternatin-3 was ∼5-fold less potent than ternatin-4 (**Figure 1––figure supplement 2**), consistent with both eEF1A binding and cell proliferation assays (Carelli et al., 2015). These data argue that didemnin and ternatin-family cyclic peptides target the proofreading stage of aa-tRNA selection to prevent aa-tRNA accommodation after GTP hydrolysis.

We substantiated eEF1A as the target of ternatin-4 by performing analogous smFRET experiments using a recombinantly expressed, human eEF1A(A399V) (**Figure 1––figure supplement 3**; Methods). The eEF1A(A399V) mutant prevents ternatin photo-affinity probe labeling of eEF1A in cells and elicits partial didemnin resistance to growth inhibition as well as nearly complete ternatin-4 resistance (Carelli et al., 2015; Krastel et al., 2015). Following a 2- minute incubation with ribosomes in the absence of inhibitor, recombinant wild-type and eEF1A(A399V) ternary complexes both promoted formation of the pre-translocation complex in which the adjacently bound P- and A-site tRNAs within the ribosome spontaneously and reversibly transit classical (∼0.7 FRET) and hybrid state positions (∼0.25-0.45 FRET; **Figure 1G**) (Budkevich et al., 2011; Ferguson et al., 2015). As expected, both didemnin and ternatin-4 effectively prevented pre-translocation complex formation by wild-type eEF1A, yielding a long-lived GA-like state (**Figure 1H**, **I**, left panels). By contrast, neither inhibitor (20 µM) had discernible effects on pre-translocation complex formation when eEF1A(A399V) was employed (**Figure 1H**, **I**, right panels). These observations further validate that the A399V mutation confers didemnin and ternatin-4 resistance during aa-tRNA selection, likely by weakening small-molecule interactions with eEF1A DIII.

### Elongation complexes trapped by didemnin and ternatin-4 exhibit distinct dynamics

As observed for drugs that target EF-Tu during bacterial aa-tRNA selection (Geggier et al., 2010; Morse et al., 2020), examination of smFRET traces obtained from pre-steady-state aa-tRNA selection studies revealed that the GA-like intermediate state captured by didemnin or ternatin-4 exhibited transient excursions to both lower- and higher-FRET states (**Figure 1C**, **D**). The apparent rates of both types of transitions were low, on the order of 0.1 s^-1^ (**Table S1**). According to the kinetic model of aa-tRNA selection defined in bacteria (Geggier et al., 2010; Morse et al., 2020), these excursions correspond to CR and AC states, respectively.

To gain insights into the mechanistic impacts of didemnin and ternatin-4 and on eEF1A, we further analyzed the individual smFRET traces using hidden Markov modeling, aiming to determine inhibitor-specific differences in the occupancy and kinetic properties of higher, transient FRET state excursions (McKinney et al., 2006; Munro et al., 2007; Qin, 2004). We focused specifically on AC-like intermediate states sampled prior to the first evidence of a fully accommodated (AC or AC-like) state, which were defined as lasting ≥150 ms (Methods).

We first assessed conformational dynamics at the ensemble level by compiling transition density plots (TDPs), in which the FRET values from each single-molecule trajectory immediately before and after a specific FRET transition are revealed as well as the relative transition frequency (McKinney et al. 2006). Comparison of TDPs in the presence of didemnin and ternatin-4 revealed that both inhibitors specifically reduced the frequency of higher-FRET transitions that normally accompany the aa-tRNA selection process (**Figure 2A-C**). Notably, excursions to AC-like states were more frequent in the presence of saturating ternatin-4 than didemnin (**Figure 2B-D**). This distinction paralleled a reduction in the overall GA-like state lifetime (**Figure 2E** and **Table S1**). These analyses therefore suggest that ternatin-4 is less efficient than didemnin in preventing conformational processes within eEF1A that allow aa-tRNA to enter the PTC.

**Figure 2.**
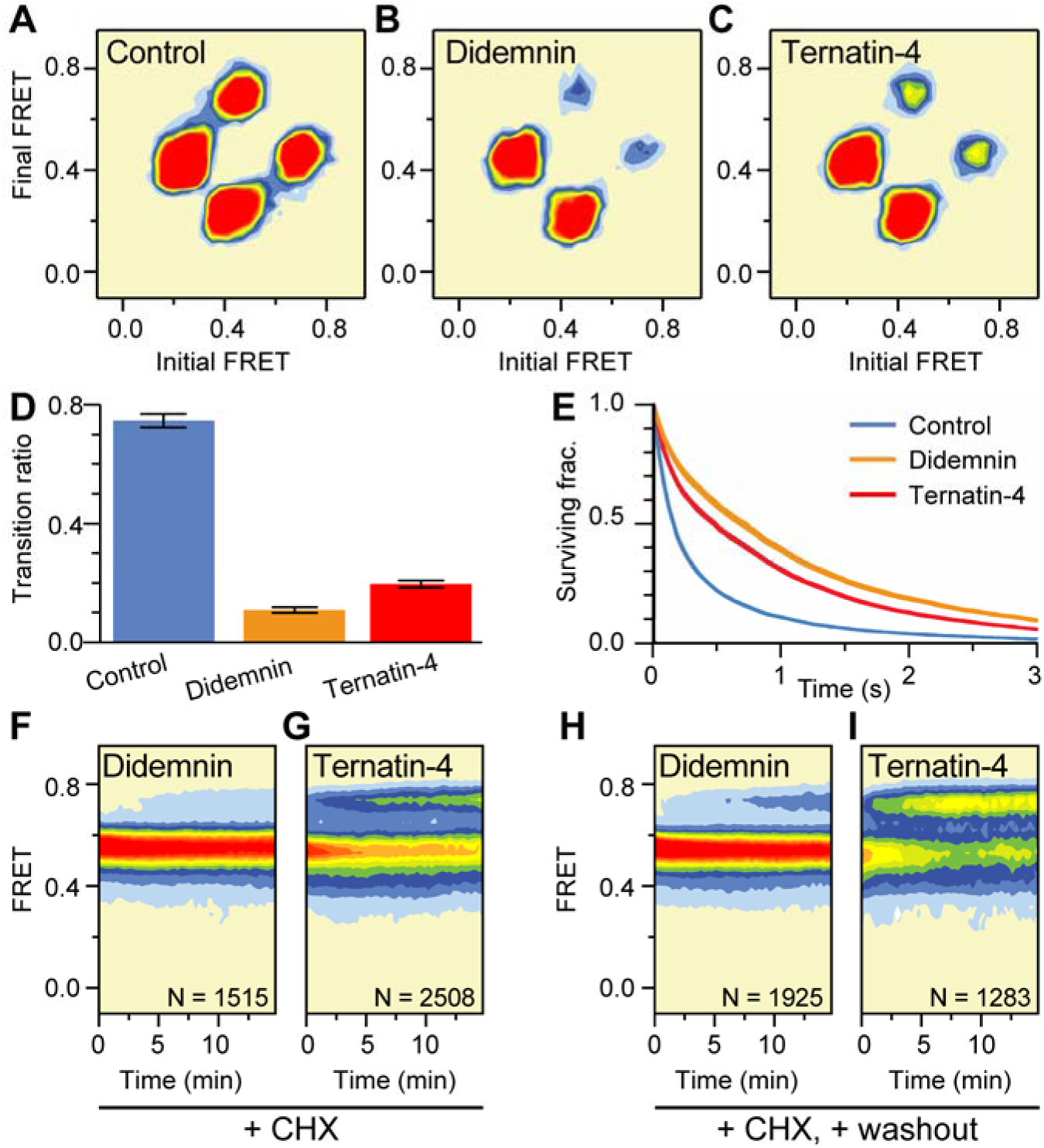
Mechanistic differences between didemnin and ternatin-4. (**A**-**C**) Transition density plots of pre-accommodated complexes reveal attenuated sampling of the high-FRET (accommodated) state comparing (A) absence of drug, or in the presence of saturating (20 µM) (B) didemnin or (C) ternatin-4. (**D**) Ratio of mid-to-high over mid-to-low transitions. Error bars: s.e.m. from 1000 bootstrap samples. (**E**) Survival plots reveal increased mid-FRET (GA) lifetimes with didemnin and ternatin-4 (line width = s.e.m. from 1000 bootstrap samples). (**F**, **G**) Population histograms of N traces after extended incubation in the presence of cycloheximide (CHX) and 20 µM (F) didemnin or (G) ternatin-4 reveals “sneak-through” to high-FRET pre-translocation complexes. (**H**) Didemnin/CHX- or (**I**) ternatin-4/CHX-stalled complexes were washed in the presence of CHX in the first second of each movie, revealing aa-tRNA accommodation, concomitant with drug dissociation. See also Figure 2—figure supplement 1.

To discern whether the excursions to AC-like states are representative of on-pathway intermediates of the aa-tRNA selection reaction coordinate, we determined the rate at which individual aa-tRNAs eventually formed pre-translocation complexes in the presence of saturating didemnin or ternatin-4. These aa-tRNA selection studies were performed at a lower frame rate (1 Hz) to reduce photobleaching and in the presence of cycloheximide (CHX, 350 µM), which depopulates hybrid tRNA positions (Ferguson et al., 2015; Garreau de Loubresse et al., 2014) that complicate analyses of the apparent didemnin and ternatin-4 stabilized, GA-like state lifetimes. Under these conditions, we observed that aa-tRNA accommodated ∼8.5-times faster in the presence of saturating ternatin-4 concentrations compared to saturating didemnin concentrations (6 × 10^-4^ s^-1^ vs. 7 × 10^-5^ s^-1^, respectively) (**Figure 2––figure supplement 1A**). These results are consistent with the model that AC-like state excursions on the human ribosome represent transient, on-pathway intermediates in the selection process. We correspondingly infer that ternatin-4 is less efficient than didemnin at inhibiting the conformational processes in eEF1A underpinning aa-tRNA accommodation during the proofreading stage of the tRNA selection.

We next sought to distinguish whether the observed excursions to AC-like states reflect differences in drug dissociation kinetics or differences in eEF1A dynamics while the drugs remain bound. To do so, we measured the rate of aa-tRNA accommodation from inhibitor-stalled GA-like states following rapid drug washout from the imaging chamber. Here, inhibitor dissociation from eEF1A is expected to enable rapid aa-tRNA accommodation, resulting in a CHX-stabilized, pre-translocation complex. Notably, the apparent drug dissociation rates were >30-fold lower than the frequency of eEF1A conformational changes that allow aa-tRNA to sample AC-like states in the presence of saturating inhibitors (**Figures 1C**, **D** and **2B**, **C**). We further conclude that ternatin-4 dissociates ∼25-fold faster than didemnin from stalled elongation complexes (∼5 × 10^-3^ s^-1^ vs. ∼2 × 10^-4^ s^-1^, respectively, **Figure 2H**, **I** and **Figure 2––figure supplement 1B-D**).

By inference from mechanistic investigations of aa-tRNA selection on the bacterial ribosome (Geggier et al., 2010; Morse et al., 2020), and the cryo-EM structure of the didemnin- stalled aa-tRNA selection intermediate isolated from rabbit reticulocyte lysate, we propose that didemnin and ternatin-4 inhibit essential domain separation processes within eEF1A after GTP hydrolysis that govern the proofreading mechanism of aa-tRNA selection. We surmise from our analyses that the FRET excursions in the presence of drug occur while didemnin and ternatin-4 remain bound to eEF1A on the ribosome. Further, we posit that AC-like excursions of aa-tRNA towards the PTC are coupled to such conformational events in eEF1A and that these events are differentially inhibited by didemnin and ternatin-4 binding.

### Ternatin-4 occupies the same binding site as didemnin on eEF1A

To compare the didemnin and ternatin-4 binding sites on eEF1A, we employed ternatin-4 in procedures analogous to those used to solve a didemnin-stalled aa-tRNA selection intermediate by cryo-EM (Shao et al., 2016). Ternatin-4 was added to rabbit reticulocyte lysate followed by immunoprecipitation procedures to pull down actively translating ribosomes via the nascent peptide (Shao et al., 2016). These efforts yielded a cryo-EM reconstruction of a ternatin-4/eEF1A/ribosome complex that resolved to 4.1 Å (**Figure 3––figure supplement 1** and **Table S2**). Global features of this map and the position of ternary complex within the ribosomal A site were highly similar to the structure stabilized by didemnin (**Figure 3A**, **B**), though the density for aa-tRNA and eEF1A was less well resolved, consistent with ternatin-4 allowing increased ternary complex mobility (**Figure 3––figure supplement 1B**).

**Figure 3.**
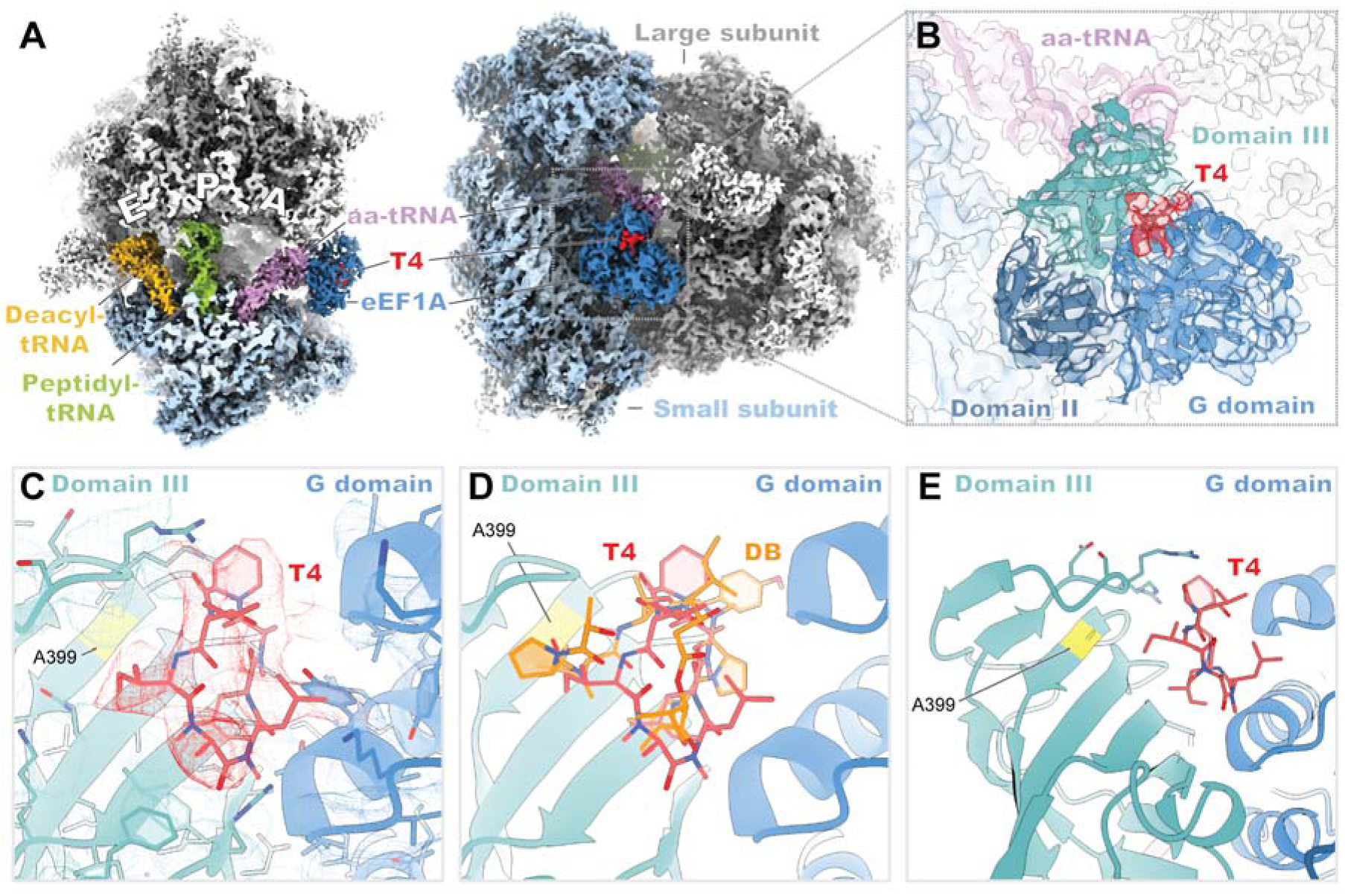
Cryo-EM structure of ternatin-4 stalled rabbit 80S-eEF1A-aa-tRNA complex. (**A**) Overview of the cryo-EM density maps of the ternatin-4-stalled rabbit elongation complex viewed from the small subunit (SSU) head domain (*left*) and into the GTPase activating center (GAC) from the leading edge (*right*) comprising the large (LSU; gray) and SSU (light blue) ribosomal subunits, peptidyl-tRNA (P site; green) and deacyl-tRNA (E site; gold), aminoacyl- tRNA in the pre-accommodated A/T state (aa-tRNA; purple), eEF1A (blue), and ternatin-4 (T4; red). (**B**) Cryo-EM density of eEF1A ternary complex on the ribosome highlighting density for T4 (red), with eEF1A colored by domain. Molecular model of eEF1A and aa-tRNA (purple) from PDB-ID: 5LZS (Shao et al., 2016) was rigid-body fit into the cryo-EM map. (**C**) Zoom-in of cryo- EM density at the interface between the G domain and domain III of eEF1A that has been assigned to T4 (red), colored as in (B). Residue A399 (yellow), which confers resistance to didemnin (DB) and T4 and when mutated to valine, is adjacent to the density for T4. (**D**) Overlay of molecular models of DB (orange) and T4 (red) from the same camera angle as (C). (**E**) View of the DB and T4 binding pocked on eEF1A, highlighting the site of a resistance mutation (A399; yellow) and the 375-391 loop of domain III between β15-16. All cryo-EM density is contoured at 3.5 σ. See also Figure 3—figure supplement 1-4.

In the presence of ternatin-4, the ribosome adopts an unrotated conformation, with the aa-tRNA in a GA-like conformation within the ribosomal A site and eEF1A bound to the GAC of the LSU, docking against the catalytic Sarcin ricin loop (**Figure 3A**). Superposition of the cryo-EM density from the ternatin-4-stalled complex with the atomic model of the didemnin-stalled complex (Shao et al., 2016) revealed clear density in the cleft between eEF1A DI (G domain) and III, which we interpret as ternatin-4 (**Figure 3B**). The position of this density overlaps with the didemnin binding site near Ala399 (**Figure 3C**-**E**), consistent with the A399V resistance mutation disrupting the drug binding pocket via steric clash. The computationally derived binding pose of ternatin on eEF1A (Sánchez-Murcia et al., 2017) also closely matched our experimental density (**Figure 3—figure supplement 2**).

To connect structural features with the inhibition mechanisms of didemnin and ternatin-4, we reconstituted human 80S ICs using reagents and procedures analogous to those used for our smFRET studies for cryo-EM analysis. Guided by our smFRET experiments, ternary complex was delivered to human ICs in the presence of either didemnin (200 nM) or ternatin-4 (20 µM) and flash frozen on cryo-EM grids within 2 minutes. The resulting human 80S tRNA selection intermediate structures were notably similar to those isolated from rabbit reticulocyte lysate with higher global nominal resolution (**Figure 3—figure supplement 3**, **4** and **Table S2**). These structures showed an increased quality of density for the tRNAs resulting from the reconstituted nature of the ICs. As was observed in the elongating rabbit structures, the cryo-EM density for eEF1A and aa-tRNA was better resolved in the didemnin-stalled human 80S structure than that stalled by ternatin-4 (**Figure 3—figure supplement 3D**, **E**). Consistent with the differences in the “on-ribosome” ternary complex dynamics revealed by smFRET, the human sample stalled by ternatin-4 required nearly three times the number of micrographs as the sample stalled by didemnin to yield a structure with well-defined cryo-EM density for eEF1A (**Figure 3—figure supplement 3**). These findings definitively show that didemnin and ternatin-4 share the same allosteric binding site on eEF1A and suggest that both inhibitors restrict conformational changes in eEF1A that accompany or facilitate aa-tRNA accommodation after GTP hydrolysis, likely related to the separation of the DI/III interface where drug binding occurs.

### Ternatin-4 traps eEF1A on the ribosome with disordered switch loops

Inspection of the G domain (DI) of eEF1A in the ternatin-4-bound rabbit 80S complex revealed no density for a γ phosphate (**Figure 4A**, **B**), consistent with ternatin-4 stalling of eEF1A after GTP hydrolysis, as was observed with didemnin (Shao et al., 2016). However, we observed differences in the stability of the switch-loops of eEF1A’s G domain when stalled by didemnin as compared to ternatin-4—elements that canonically become disordered following GTP hydrolysis and P_i_ release in Ras-family GTPases (Bourne et al., 1991; Gasper and Wittinghofer, 2019). In the ternatin-4-bound rabbit 80S complex, the switch-I and -II elements of eEF1A displayed particularly weak cryo-EM density (**Figure 4A**, **B**). By contrast, the previously determined cryo-EM structure of the didemnin-trapped rabbit 80S ribosome (Shao et al., 2016) exhibited comparatively ordered cryo-EM density for both switch loops despite P_i_ dissociation (**Figure 4B**). This was also true of the didemnin-bound human 80S ribosome reported here (**Figure 4—figure supplement 1A**, **B**).

**Figure 4.**
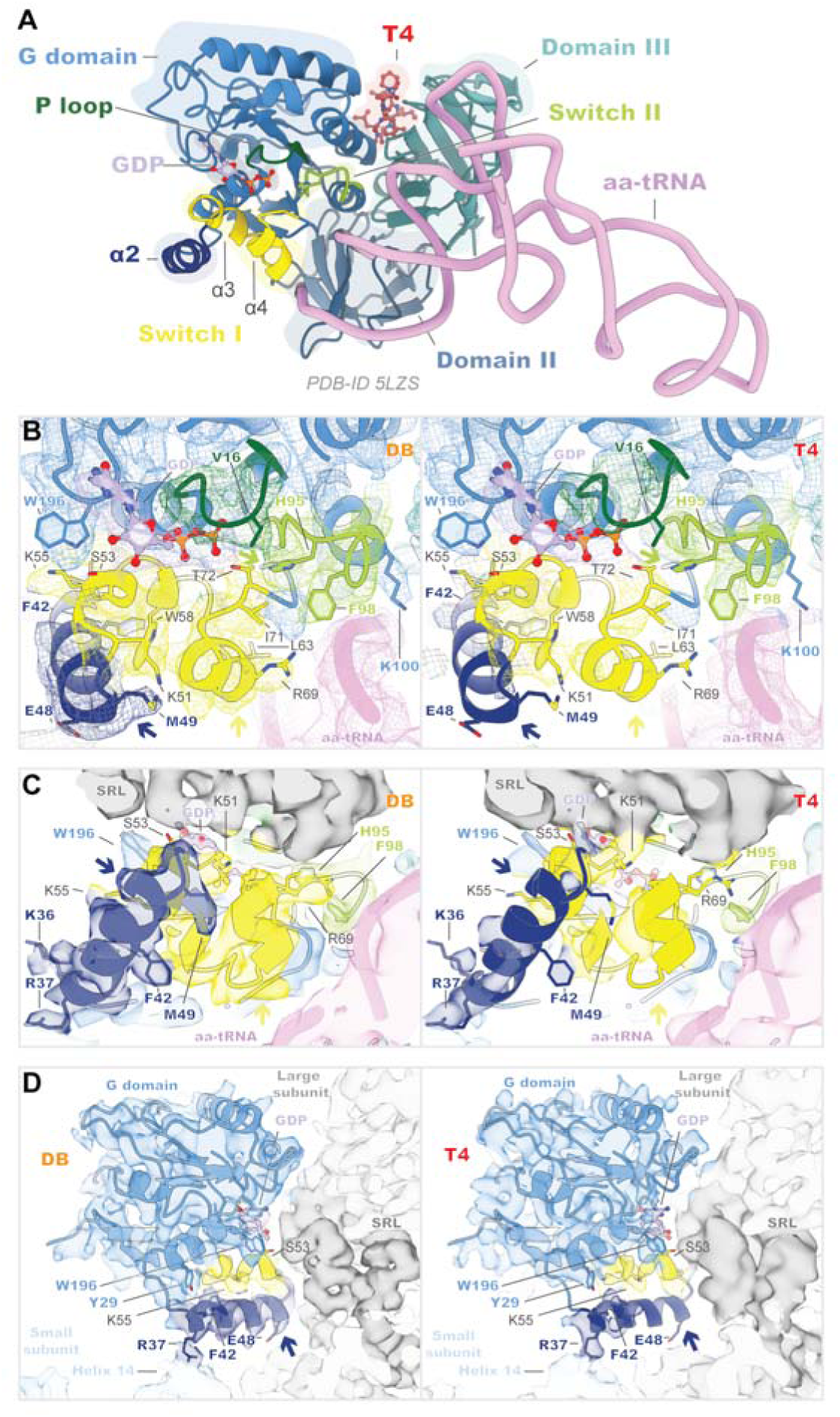
eEF1A is more dynamic when bound to ternatin-4 than to didemnin. (**A**) Overview of the domain architecture of the eEF1A ternary complex from PDB-ID: 5LZS (Shao et al., 2016) bound to ternatin-4 (T4; red) as viewed from the leading edge of the rabbit 80S ribosome and the Sarcin ricin loop (SRL). G-domain (blue) elements include switch I (yellow), switch II (lime), the P loop (dark green), helix α2 (dark blue), and a bound GDP (light purple) in the nucleotide binding pocket. (**B**-**D**) Molecular models from PDB-ID: 5LZS (Shao et al., 2016) were rigid-body fit into the cryo-EM density shown in (B) mesh and (C, D) surface representation of the eEF1A G domain when stalled with didemnin (DB; orange; EMD-4130; *left*) (Shao et al., 2016) or T4 (*right*) on the elongating rabbit 80S ribosome, colored as in (A). Colored arrows indicate regions of weakened cryo-EM density in the T4-stalled eEF1A G domain in the C terminus of helix α2 (dark blue), the C terminus of switch I (yellow) and the catalytic His95 of switch II (lime). Panels highlight (B) the nucleotide binding pocket of eEF1A and switch loop architecture, (C) the interface between eEF1A and the SRL (dark gray), and (D) the junction between the SRL, eEF1A helix α2, and small subunit (SSU; light blue) rRNA helix 14. All cryo-EM density is contoured at 3 σ. See also Figure 4—figure supplement 1.

Comparatively, the ternatin-4-stalled structures displayed weakened density for the putative catalytic His95 of switch II, for the hydrophobic gate elements of the p loop (Val16) and switch I (Ise71), and for the C-terminal helix (α4) of switch I (**Figure 4B** and **Figure 4—figure supplement 1B**). Further, we observed a loss of α4 residue Arg69 contact with the aa-tRNA minor groove (**Figure 4B** and **Figure 4—figure supplement 1B**), potentially contributing to the increase in aa-tRNA dynamics of ternatin-4-stalled ternary complexes seen by smFRET (**Figure 2A**-**C**). We did, however, observe maintenance of switch-II hydrophobic gate residue Phe98 and possibly strengthened contact between switch-I residue Lys51 and the Sarcin ricin loop (**Figure 4C** and **Figure 4—figure supplement 1C**). We speculate that these interactions may be critical for stabilizing eEF1A binding to the LSU and for preventing aa-tRNA accommodation.

Notably, we observed weakened density for the C-terminal portion of helix α2 of eEF1A in the ternatin-4-stalled complexes (**Figure 4C** and **Figure 4—figure supplement 1C**). This helical insertion, which is not present in the bacterial equivalent of eEF1A, makes direct contact with both the SSU and LSU and appears to be stabilized by the active conformation of switch I in the rabbit didemnin-stalled 80S complex (Shao et al., 2016) as well in human (**Figure 4D** and **Figure 4—figure supplement 1D**). In the rabbit ternatin-4-stalled structure, contact between α2 and the LSU appeared weakened compared to the didemnin-stalled structure, while contact between the α2 - between the didemnin- and ternatin-4-stalled complexes in this region were less pronounced (**Figure 4—figure supplement 1D**). In fact, this α2 to SSU contact appeared intact in both human structures, bolstered by strong α2 Phe42 contact with the switch-I N terminus (**Figure 4C**). This may indicate that loss of α2 contact with the LSU precedes dissociation from the SSU along the tRNA selection reaction coordinate.

We infer from the observed disordering of aa-tRNA and eEF1A in the ternatin-4-stalled complexes likely reflects post-GTP hydrolysis eEF1A dynamics within the ternary complex. Similar concepts were put forward through structural studies of the bacterial aa-tRNA selection inhibitor kirromycin (Fischer et al., 2015; Schmeing et al., 2009), which targets the equivalent EF-Tu binding pocket as didemnin and ternatin-4. We note in this context that the increased dynamics evidenced by the cryo-EM structures of ternatin-4-stalled complexes correlated with those evidenced by smFRET within the drug-stalled, GA-like state (**Figures 1**, **2**).

### Didemnin, but not ternatin-4, irreversibly inhibits protein synthesis in cells

The observed differences in the structure and dynamics of didemnin- and ternatin-4-stalled eEF1A on the ribosome prompted us to investigate whether similar kinetic differences could be discerned in cells. After 4 hours of continuous treatment, both compounds potently inhibited protein synthesis in HCT116 cells (**Figure 5A**, didemnin IC_50_ ∼7 nM; ternatin-4 IC_50_ ∼36 nM), as measured by metabolic labeling with homopropargylglycine (Hpg) and flow cytometry analysis (Beatty et al., 2006). Cells treated with ternatin-4 (500 nM, ∼14× IC_50_), followed by rigorous washout, recovered protein synthesis rates to ∼25% of starting levels within 22 hours (**Figure 5B**, **C**). By contrast, protein synthesis was undetectable for at least 22 hours followed by washout in cells treated with saturating didemnin (100 nM, ∼14× IC_50_).

**Figure 5.**
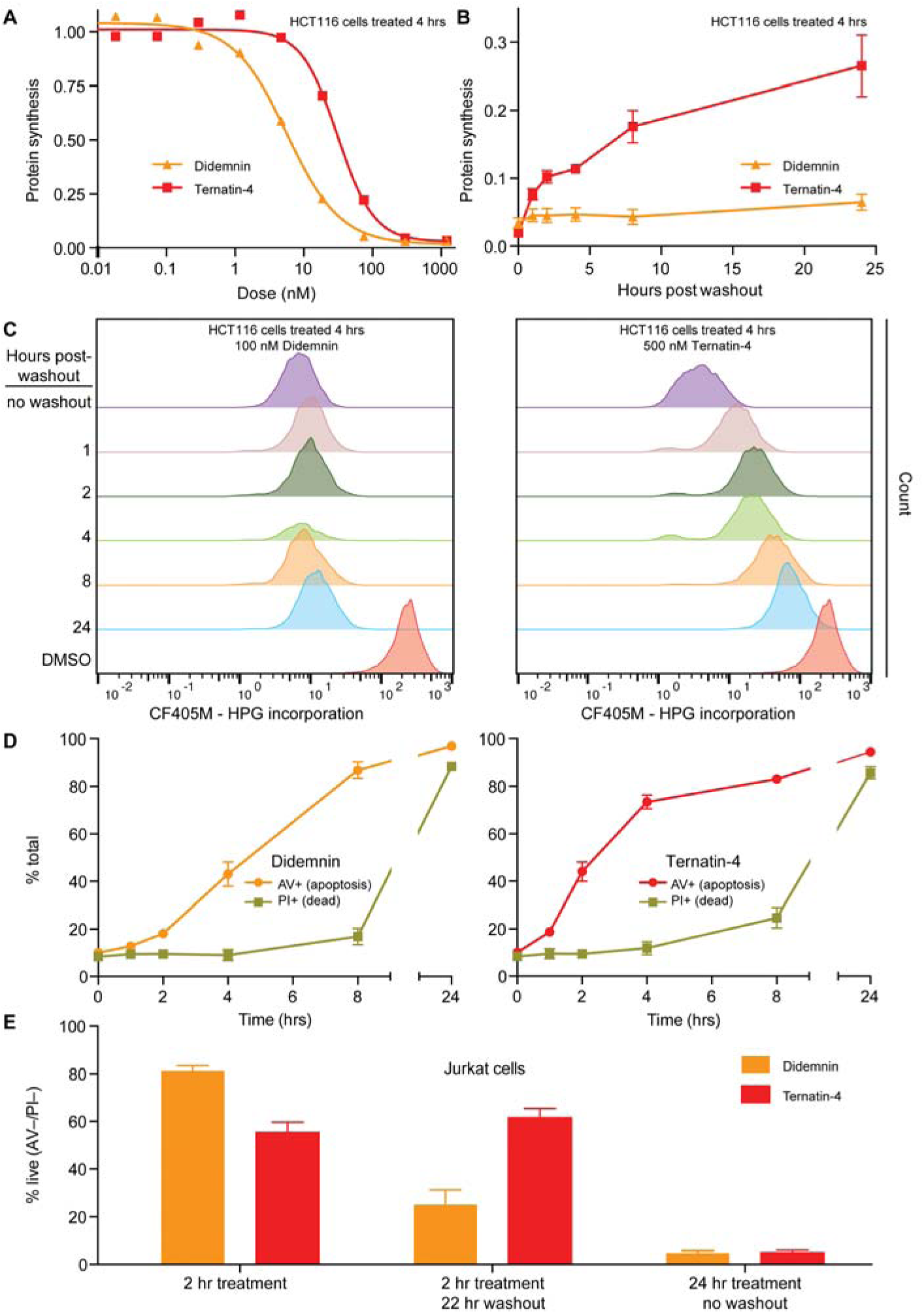
Cellular effects of ternatin-4, but not didemnin, are reversible upon washout. (**A**) Dose-dependent effects of didemnin (orange) and ternatin-4 (red) on protein synthesis in HCT116 cells under continuous treatment (4 hours). Protein synthesis was quantified by homopropargylglycine pulse (1 hr) followed by fixation and copper-mediated conjugation to CF405M azide fluorophore and analyzed by FACS. (**B**) HCT116 cells were treated with didemnin (100 nM; orange) or ternatin-4 (500 nM, red) for 4 hours, followed by washout. Protein synthesis was quantified as in (A) at 1, 2-, 4-, 8-, or 24-hours post-washout. (**C**) Histograms corresponding to panel (B) for didemnin (*left*) and ternatin-4 (*right*). (**D**) Jurkat cells were treated with didemnin (100 nM, *left*) or ternatin-4 (500 nM; *right*) or for 1, 2, 4, 8, or 24 hours, stained with Annexin V-FITC (AV) and propidium iodide (PI; green), and analyzed by FACS. (**E**) Jurkat cells were treated with didemnin (100 nM; orange) or ternatin-4 (500 nM; red) for 2 hours followed by washout and 22-hour incubation in drug-free media or for 24 hours and analyzed as in (D). See also Figure 5—figure supplement 1.

To substantiate these kinetic differences, we monitored time-dependent induction of apoptosis under conditions of continuous drug exposure or after a brief pulse, followed by washout. We used Jurkat cells which are known to undergo rapid apoptosis in the presence of didemnin (Baker et al., 2002). Continuous exposure to didemnin or ternatin-4 for 24 hours induced apoptosis in >95% of the cells, with didemnin being ∼7-fold more potent than ternatin-4 (**Figure 5––figure supplement 1**, didemnin IC_50_ ∼4 nM; ternatin-4 IC_50_ ∼30 nM). Treatment with saturating didemnin or ternatin-4 induced membrane phosphatidylserine exposure (an early marker of apoptosis) within 2-4 hours in a subpopulation of cells (**Figure 5D**). A 2-hour pulse with saturating ternatin-4 was sufficient to induce apoptosis in ∼40% of the cells, whereas rigorous washout followed by 22-hour incubation in compound-free media resulted in no further cell death (**Figure 5E**). By contrast, cell death increased from ∼20% after a 2-hour didemnin pulse to ∼75% after drug washout, consistent with didemnin’s ability to inhibit protein synthesis in a sustained, washout-resistant manner (**Figure 5C**). Collectively, these results demonstrate clear differences in cellular pharmacology between didemnin and ternatin-4 under conditions of transient drug exposure followed by washout, which correlate with the ∼25-fold higher dissociation rate of ternatin-4 observed by smFRET (**Figure 2**).

## Discussion

In this study, we combined the complementary methods of smFRET, cryo-EM, and *in vivo* translation measurements to reveal the molecular mechanisms by which didemnin and ternatin- 4 inhibit translation in mammals. Both compounds bind an allosteric site at the eEF1A DI/III interface, which likely prevents the inter-domain rearrangements that allow for aa-tRNA accommodation into the ribosomal A site and eEF1A dissociation from the ribosome. As a result, didemnin and ternatin-4 trap eEF1A(GDP)aa-tRNA ternary complex for extended periods during an intermediate stage of the aa-tRNA selection process. Compared to didemnin, ternatin- 4 traps eEF1A in a more dynamic state, characterized by conformational heterogeneities and a reduced extent of observable eEF1A-ribosome contacts. While didemnin and ternatin-4 have similar biochemical potencies in *in vitro* translation reactions, our findings are consistent with ternatin-4 dissociating ∼25× more rapidly than didemnin under washout conditions. Correspondingly, while didemnin and ternatin-4 inhibited protein synthesis and induced apoptosis at similar concentrations, didemnin exhibited quasi-irreversible cellular effects whereas the effects of ternatin-4 were reversible on washout.

The *in vivo* residence time of didemnin and ternatin-4 may influence their efficacy and toxicity profiles. Given the rapid response of some cell lines to eEF1A inhibitors, there is a possibility for achieving efficacy against rapidly proliferating cells *in vivo* with a ternatin-like inhibitor, while sparing the broad toxicity of irreversible translation inhibition. Indeed, we have recently found that a hydroxylated variant of ternatin-4 is efficacious in a mouse model of MYC-dependent B cell lymphoma (Wang et al., 2020). Additionally, subtle differences in the mechanism of competitive inhibitors can lead to dramatic differences in toxicity and potential therapeutic utility, as was observed for vinca domain-binding tubulin modulators (Wieczorek et al., 2016). Recent work has found ternatin-4 to have an IC_90_ of 15 nM against SARS-CoV-2 *in vitro* (Gordon et al., 2020), underscoring the value of translation-targeting drugs to human therapeutics. A related study found that plitidepsin (also known as dehydrodidemnin B, a close structural relative of didemnin B originally advanced in clinical trials for the treatment of multiple myeloma) also possessed potent antiviral activity against SARS-CoV-2 with an IC_90_ of 0.88 nM (White et al., 2021). Identifying the molecular context that determines sensitivity or resistance to didemnins and ternatins will likely be invaluable for identifying unexploited and forthcoming therapeutic applications of eEF1A inhibitors.

## Materials and Methods

### Ribosome subunit isolation for human smFRET and cryo-EM analysis

Human ribosome subunits were isolated from adherent HEK293T cells as described previously (Ferguson et al., 2015). Cells were grown in high-glucose Dulbecco’s modified Eagle’s medium (Life Technologies) supplemented with 10% fetal bovine serum (Atlanta Biologicals) and 1% penicillin/streptomycin (Life Technologies). At ∼75% confluency, 350 µM cycloheximide (CHX) was added to the medium and incubated for 30 min. Cells were detached with 0.05% Trypsin-EDTA supplemented with 350 µM CHX, pelleted and flash-frozen in liquid nitrogen for storage.

For ribosome isolation, cells were thawed on ice and resuspended in lysis buffer (20 mM Tris HCl pH 7.5, 10 mM KCl, 5 mM MgCl_2_, 1 mM DTT, 5 mM putrescine, 350 µM CHX, 4 U/ml RNAse Out (Thermo Fisher), 1 × Halt Protease Inhibitor Cocktail without EDTA (Thermo Fisher)), followed by addition of 0.5% (v/v) NP-40, 0.5% (w/v) sodium deoxycholate, and 20 U/ml Turbo DNAse (ThermoFisher) and incubation at 4°C for 20 minutes while rotating. Lysate was clarified by brief centrifugation, loaded onto pre-chilled 10-50% sucrose gradients prepared in polysome gradient buffer (20 mM Tris HCl pH 7.5, 10 mM KCl, 5 mM MgCl_2_, 1 mM DTT, 5 mM putrescine, 350 µM CHX), and centrifuged for 3 hours at 110 krcf, 4°C. Gradients were analyzed following standard procedures on a gradient fractionator (Brandel) with UV absorbance detector (Teledyne Isco), polysome fractions were collected and pelleted for 18 hours at 125 krcf, 4°C. Polysome pellets were rinsed and resuspended in resuspension buffer (20 mM Tris HCl pH 7.5, 50 mM KCl, 1.5 mM MgCl_2_, 1 mM DTT), followed by addition of 1 mM puromycin, raising the KCl concentration to 500 mM, 30 minutes incubation at 4°C with rotation, and 15 minutes incubation at 37°C. The solution was cleared by centrifugation, loaded onto 15-30% sucrose gradients prepared in subunit gradient buffer (5 mM Tris HCl pH 7.5, 500 mM KCl, 2.5 mM MgCl_2_, 1 mM DTT) and centrifuged for 14 hours at 50 krcf, 20°C. Gradients were analyzed as before and individual 40S and 60S subunit fractions were collected, followed by centrifugation for 3 hours (60S) or 6 hours (40S) at 425 krcf, 4°C. Subunit pellets were resuspended in 80S polymix buffer (30 mM HEPES pH 7.5, 5 mM MgCl_2_, 50 mM NH_4_Cl, 5 mM Putrescine, 2 mM Spermidine, 1 mM DTT), aliquoted, and flash-frozen in liquid nitrogen.

### Human 80S IC formation for smFRET and cryo-EM

For smFRET analyses, synthetic mRNA (Dharmacon, sequence CAA CCU AAA ACU UAC ACA CCC UUA GAG GGA CAA UCG **AUG UUC AAA** GUC UUC AAA GUC AUC) was prepared for surface immobilization in the following way: 45 µM each of DNA oligonucleotides A (AAA AAA AAA AAA AAA AAA AAA AAA AAA AAA) and B (GTA AGT TTT AGG TTG CCC CCC TTT TTT TTT TTT TTT TTT TTT TTT TTT TTT with 3’-Biotin-TEG modification) in hybridization buffer (10 mM HEPES pH 7, 150 mM KCl, 0.5 mM EDTA) were heated to 95°C for 5 minutes and annealed on ice for 5 min. The annealed oligonucleotide (20 µM) and mRNA (20 µM) in hybridization buffer were incubated for 5 minutes at 37°C and for 5 minutes on ice. For smFRET analyses, tRNA^fMet^ isolated from *E. coli* was labeled with Cy3 at the s^4^U8 residue following established procedures (Blanchard et al., 2004b). Cy3-labled (smFRET) or unlabeled (cryo-EM) tRNA^fMet^ was aminoacylated by incubation of 30 or 120 pmol of tRNA, respectively, with 50 mM Tris pH 8, 25 mM KCl, 100 mM NH_4_Cl, 10 mM MgCl_2_, 1 mM DTT, 5 mM ATP, 0.5 mM EDTA, 5 mM Met amino acid, 600 nM Met-RS in a 10-20 µl reaction at 37°C for 15 min.

Human 80S ICs for smFRET and cryo-EM were formed by incubating 20 or 100 pmol of 40S subunits (heat-activated at 40°C for 5 min), respectively, and 4× molar excess of prepared mRNA/DNA duplex (smFRET) or mRNA (cryo-EM) in 80S polymix for 10 minutes at 37°C and for 5 minutes on ice. The freshly prepared aminoacylated tRNA was added, followed by incubation for 10 minutes at 37°C and for 5 minutes on ice. Equimolar 60S subunits (heat-activated at 40°C for 5 min) were added (final reaction volume 50-100 µl) and the reaction mixture was incubated for 20 minutes at 37°C and for 5 minutes on ice. MgCl_2_ concentration was then adjusted to 15 mM and the reaction mixture was loaded onto a 10-30% sucrose gradient prepared in 80S polymix as above but with 15 mM MgCl_2_. Gradients were centrifuged at 150 krcf, 4°C for 90 minutes and analyzed as described above, collecting the fraction corresponding to 80S complexes. For smFRET, the resulting fractions aliquoted and flash-frozen in liquid nitrogen. For cryo-EM analysis, 80S ICs were pelleted at 150 krcf, 4°C for 90 minutes and resuspended in a minimal volume of 80S polymix with 5 mM MgCl_2_ for a final concentration of 2-3 µM.

### Purification of rabbit eEF1A

Rabbit reticulocyte lysate (Green Hectares) was thawed, supplemented with 1 × Mammalian ProteaseArrest (G-Biosciences), 1 mM phenylmethane sulfonyl fluoride, and 500 mM KCl, layered onto a cushion of 20 mM Tris HCl pH 7.5, 1 M sucrose, 500 mM KCl, 5 mM MgCl_2_, 1 mM DTT, 20% glycerol, and centrifuged for 14 hours at 125 krcf, 4°C. The supernatant was fractionated by precipitation with increasing concentrations of (NH_4_)_2_SO_4_ by gradual addition of saturated solution while stirring at 4°C. The fraction corresponding to 30-40% saturation was centrifuged for 20 minutes at 25 krcf, 4°C. Pellets were resuspended into and dialyzed against Buffer A (20 mM Tris-HCl pH 7.5, 50 mM KCl, 0.1 mM EDTA, 0.25 mM DTT, 20% glycerol).

The sample was then further purified by three ion-exchange steps. In each step, fractions enriched in eEF1A were detected by Western Blot analysis (primary antibody: Millipore 05-235, used at 1:2000 dilution in TBST with 5% w/v dry milk). The columns used for the three steps were (i) DEAE FF HiPrep 16/10, (ii) SP HP 5 mL HiTrap, (iii) Mono S 5/50 GL (all from GE). In all three steps, Buffer A was used for column equilibration and sample loading; a gradient into Buffer B (identical to Buffer A but with 1 M KCl) was used for elution. In step (i), eEF1A was enriched in the flow-through, in subsequent steps, it was enriched in the eluted fractions. After steps (i) and (ii), the most enriched fraction as identified by Western Blot was dialyzed against Buffer A; after step (iii), the final product (∼85% pure) was dialyzed against storage buffer (20 mM Tris HCl pH 7.5, 25 mM KCl, 6 mM BME, 5 mM Mg(OAc)_2_, 60% glycerol) and stored at -20°C.

### smFRET analysis of aa-tRNA selection

smFRET experiments were performed on a custom-built prism-type TIRF microscope as described previously (Juette et al., 2016). Briefly, biotinylated 80S ICs were immobilized in flow cells treated with a mixture of polyethylene glycol (PEG) and PEG-biotin and functionalized with streptavidin (Blanchard et al., 2004b). For aa-tRNA selection experiments, ternary complex containing *E. coli* Phe-tRNA^Phe^ labeled with Cy3 at position acp^3^U47 (Blanchard et al., 2004b), rabbit eEF1A, and GTP was delivered by manual injection at a final concentration of 20 nM. All experiments were performed in 80S polymix buffer with 5 mM MgCl_2_ as above. A 532 nm diode-pumped solid-state laser (Opus, LaserQuantum) was used for Cy3 excitation, fluorescence was collected through a 60×/1.27 NA water-immersion objective (Nikon), spectrally separated using a T635lpxr-UF2 dichroic mirror (Chroma) and imaged onto two cameras (Orca-Flash 4.0 v2, Hamamatsu). Time resolution was 15 ms for all experiments except for the sneak- through/wash-out experiments shown in Figure 2F-I, which were performed at 1 s time resolution.

### smFRET data processing and analysis

Single-molecule fluorescence and FRET traces were extracted and further analyzed using our freely available MATLAB-based software platform SPARTAN (Juette et al., 2016) (http://scottcblanchardlab.com/software), extended with custom scripts. For display of example traces (Figure 1) and all quantitative analysis, traces were idealized using the sequential k-means algorithm based on a hidden Markov model (Qin, 2004). For display of non-equilibrium aa-tRNA selection data, all detected events were post-synchronized by aligning them to the first appearance of FRET. FRET contour plots were generated by compiling 2-dimensional histograms of FRET occupancy over time for all traces. For dose-response curves (Figure 1E, F and Figure 1—figure supplement 1), accommodated molecules were defined as events spending 300 ms or more in high-FRET. EC_50_ values were obtained by fitting a Hill equation to the accommodated fraction as a function of drug concentration. To assess kinetic differences during aa-tRNA selection (Figure 2), the analysis of each idealized FRET trace was restricted to the time interval prior to the first dwell in high FRET lasting 150 ms or more (shorter than the above-mentioned definition of accommodated molecules to reduce contributions from molecules achieving hybrid states after accommodation). These truncated, idealized traces were used for the computation of state lifetimes and transition ratios. Error bars in Figure 1E, F, Figure 1— figure supplement 1, and Figure 2D, E represent standard errors obtained by bootstrap analysis (1000 samples) of the pooled data from all experimental repeats.

### Rabbit in vitro translation and cryo-EM sample preparation

*In vitro* translation reactions of a transcript encoding 3× Flag-tagged KRas were performed in a rabbit reticulocyte lysate (RRL) system at 32°C as previously described (Shao et al., 2016; Sharma et al., 2010). A transcript encoding 3× Flag-tagged KRas was translated *in vitro*. A final concentration of 50 µM ternatin-4 was added after 7 minutes to stall ribosome-nascent chain complexes (RNCs) at the stage of aa-tRNA delivery by eEF1A and the reaction allowed to proceed to 25 min. A 4 ml translation reaction was directly incubated with 100 μl (packed volume) of anti-Flag M2 beads (Sigma) for 1 hour at 4°C with gentle mixing. The beads were washed sequentially with 6 ml of buffer (50 mM HEPES pH 7.4, 5 mM Mg(OAc)_2_, and 1 mM DTT) containing the additional components as follows: (1) 100 mM KOAc and 0.1% Triton X- 100; (2) 250 mM KOAc 0.5% and Triton X-100; (3, RNC buffer) 100 mM KOAc. Two sequential elutions were carried out with 100 μl 0.1 mg/ml 3× Flag peptide (Sigma) in RNC buffer at room temperature for 25 min. The elutions were combined and centrifuged at 100,000 rpm at 4°C for 40 minutes in a TLA120.2 rotor (Beckman Coulter) before resuspension of the ribosomal pellet in RNC buffer containing 1 μM ternatin-4. The resuspended RNCs were adjusted to 120 nM and directly frozen to grids for cryo-EM analysis.

R2/2 grids (Quantifoil) were covered with a thin layer of continuous carbon (estimated to be 50 Å thick) and glow discharged to increase hydrophilicity. The grids were transferred to a Vitrobot MKIII (FEI) with the chamber set at 4°C and 100% ambient humidity. Aliquots of purified RNCs (3 μl, ∼120 nM concentration in 50 mM HEPES pH 7.4, 100 mM KOAc, 5 mM Mg(OAc)_2_, 1 mM DTT and 1 μM ternatin-4) were applied to the grid and incubated for 30 s, before blotting for 3 s to remove excess solution, and vitrified in liquid ethane.

### Sample preparation for human cryo-EM structure determination

Gold R1.2/1.3 300 mesh grids (UltrAuFoil) were plasma cleaned (ArO_2_, 7 s) and transferred to a Vitrobot MKII (FEI) with the chamber set at 4°C and 100% ambient humidity. Aliquots of purified human 80S ICs in 80S polymix buffer were thawed and brought to 0.2 μM didemnin B (didemnin) or 20 μM ternatin-4. Ternary complex containing *E. coli* Phe-tRNA^Phe^, rabbit eEF1A, GTP, and either 0.2 μM didemnin or 20 μM ternatin-4 were added to 80S ICs for final concentrations of ∼200 nM 80S and aa-tRNA. aa-tRNA selection reactions were applied to the grid (3 μl) and incubated for ∼45 s, before blotting for 2-3 s to remove excess solution, and vitrified in liquid ethane.

### Cryo-EM data collection and image processing for the rabbit structure

All micrographs of rabbit 80S ribosomes were taken on an FEI Titan Krios microscope (300 kV) equipped with an FEI Falcon II direct-electron detector using quasi-automated data collection (EPU software, FEI). Movies were recorded at a magnification of ∼135,000 ×, which corresponds to the calibrated pixel size of 1.04 Å per pixel at the specimen level. During the 1-s exposure, 40 frames (0.06 s per frame) were collected with a total dose of around 40 e^−^ per Å^2^. Movie frames were aligned using whole-image motion correction (Li et al., 2013). Parameters of the contrast transfer function (CTF) for each motion-corrected micrograph were obtained using Gctf (Zhang, 2016). Visual inspection of the micrographs and their corresponding Fourier transforms was used to removed micrographs due to astigmatism, charging, contamination, and/or poor contrast.

Ribosome particles were selected from the remaining micrographs using semi-automated particle picking implemented in RELION 1.4 (Scheres, 2015). Reference-free two-dimensional class averaging was used to discard non-ribosomal particles. The retained particles underwent an initial three-dimensional refinement using a 30 Å low-pass filtered cryo-EM reconstruction of the mammalian ribosomal elongation complex (EMDB-4130) as an initial model. After refinement, the particles were then subjected to three-dimensional classification to separate different compositions and conformations of the ribosome complexes and isolate particles with high occupancy of the desired factors. From this classification, particles containing P- and E-site tRNAs were selected and re-refined. The movement of each particle within this subset was further corrected using RELION 1.4 (Scheres, 2015). The resulting ‘shiny’ particles were subjected to focused classification with signal subtraction (FCwSS) (Bai et al., 2015) to isolate particles containing pre-accommodated aa-tRNA and eEF1A. An additional round of 3D refinement was used to obtain the final map, which reached an overall resolution of 4.1 Å based on the Fourier shell correlation (FSC) 0.143 criterion (Rosenthal and Henderson, 2003). During post-processing, high-resolution noise substitution was used to correct for the effects of a soft mask on FSC curves (Chen et al., 2013) and density maps were corrected for the modulation transfer function (MTF) of the Falcon II detector and sharpened by applying a negative B-factor that was estimated using automated procedures (Rosenthal and Henderson, 2003). See Figure 3—figure supplement 1 and Table S2 for details.

### Cryo-EM data collection and image processing for the human structures

All micrographs of human 80S ribosomes were taken on an FEI Titan Krios microscope (300 kV) equipped with an Gatan K2 Summit direct electron detector using Leginon MSI (Suloway et al., 2005) data collection. For didemnin, movies were recorded in counting mode at a magnification of 105,000 × (∼1.072 Å^2^ per pixel) with 10-s exposure for 50 frames (0.2 s per frame) with a total dose of around 67 e^−^ per Å^2^. For ternatin-4, movies were recorded in super resolution mode at a magnification of 105,000 × (∼1.096 Å^2^ per pixel after 2×binning) with 10-s exposure for 50 frames (0.2 s per frame) with a total dose of around 70 e^−^ per Å^2^. See Table S2 for details. Movie frames were aligned using whole-image motion correction (Li et al., 2013). CTF parameters of the contrast transfer function for each motion-corrected micrograph were obtained using CTFFIND4 (Rohou and Grigorieff, 2015). Visual inspection of the micrographs and their corresponding Fourier transforms was used to removed micrographs due to astigmatism, charging, contamination, and/or poor contrast.

Ribosome particles were selected from the remaining micrographs using semi-automated particle picking implemented in RELION 2.0 (Kimanius et al., 2016). Reference-free two-dimensional class averaging was used to discard non-ribosomal particles. The remaining particles were subjected to two rounds of three-dimensional classification with alignment to separate different compositions and conformations of the ribosome complexes, sorting first for 80S particles followed by sorting for rotated and unrotated small ribosomal subunits. Particles with unrotated subunits were subjected to focused classification and FCwSS (Bai et al., 2015) to isolate particles containing pre-accommodated aa-tRNA and eEF1A. An additional round of 3D refinement was used to obtain the final maps, which reached overall resolutions of 3.2 Å and 3.8 Å for didemnin and ternatin-4, respectively, based on the FSC 0.143 criterion (Rosenthal and Henderson, 2003). During post-processing, noise substitution was used to correct for the effects of a mask on FSC curves (Chen et al., 2013) and density maps were corrected for the MTF of the K2 detector and sharpened by applying a -20 B-factor and a 4 Å low pass filter (Rosenthal and Henderson, 2003). See Figure 3—figure supplement 3 and Table S2 for details.

### Cryo-EM map interpretation

Density map values were normalized to mean = 0 and standard deviation (σ) = 1 in UCSF Chimera (Pettersen et al., 2004) using the vop scale function. The pixel size of each map was calibrated against a 2.9 Å resolution structure of the mammalian ribosome (PDB-ID: 6QZP) (Natchiar et al., 2017) and maps were aligned on the 60S core. The model of the mammalian ribosomal elongation complex trapped with didemnin (PDB-ID: 5LZS) (Shao et al. 2016) was docked into the cryo-EM maps with Chimera (Pettersen et al., 2004). Density present in the didemnin binding site was interpreted as belonging to didemnin or ternatin-4, respectively. A model for ternatin-4 was rigid-body fit into the cryo-EM density for figure images. However, as the density was insufficiently resolved to unambiguously place a model for ternatin-4, the density was left unmodelled.

### Homoprogargyl glycine metabolic labeling

HCT116 cells were seeded in 24 well plates at 30,000 cells/well and incubated overnight before 4-hour treatment with compound. For experiments involving washout, cells were washed twice with 1 ml complete media, followed by alternating quick and 5 minutes 37°C washouts repeated 4 times each (O’Hare et al., 2013). After appropriate incubations, cells were washed once with phosphate-buffered saline (PBS), then exchanged to methionine- and cysteine-free DMEM (Gibco) supplemented with 10% dialyzed FBS (Sigma), glutamine (2 mM), cysteine (2 mM), homopropargyl glycine (1 mM; Kerafast), and appropriate drug for 1 hour. Media was then aspirated, cells were trypsinized and transferred to 96-well plates, washed once with PBS, and fixable live/dead stained with Zombie Red amine-reactive dye (BioLegend) according to the manufacturer’s instructions. Cells were fixed in 2% paraformaldehyde in PBS for 10 minutes at room temperature, and then permeabilized in PBS supplemented with 0.1% saponin and 3% FBS. Samples in 25 µl permeabilization buffer were subjected to copper-catalyzed alkyne-azide conjugation to CF405M-azide (Biotium) by addition of 100 µl click reaction mix (50 mM HEPES pH 7.5, 150 mM NaCl, 400 µM TCEP, 250 µM TBTA, 200 µM CuSO_4_, 5 µM azide). After overnight incubation at room temperature in the dark, samples were washed 3× with permeabilization buffer, 2× FACS buffer (PBS –Mg/Ca + 2% FBS + 2 mM EDTA) and analyzed by flow cytometry (MACSQuant VYB). Data analysis was performed with FlowJo software (Tree Star). Dead cells (Zombie Red +) were excluded from analysis (typically representing <15% of total cells), leaving at least 500 live cells for each data point, but additional cells were analyzed when possible (up to 10,000). IC_50_ values and plotted data points (Figure 5) are given as the mean of three independent determinations ± standard error.

### Apoptosis assay by annexin V/propidium iodide staining

Jurkat cells at 0.5 × 10^6^ cells/ml were treated with compound as indicated. For experiments involving washout, cells were washed twice with 1 ml complete media by pelleting cells for 3 minutes at 1.3k × *g* and aspirating media, followed by alternating quick and 5 minutes 37°C washouts repeated 4 times each (O’Hare et al., 2013). Cells were stained with annexin V-FITC and propidium iodide (BD Pharmingen) according to the manufacturer’s instructions and analyzed by flow cytometry (MACSQuant VYB). The mean of three independent experiments ± standard error is plotted. For each drug treatment condition, 10,000 cells were analyzed.

### Flag-eEF1A purification

HCT116 cells stably expressing Flag-eEF1A1(WT or A399V)-P2A-mCherry (Carelli et al., 2015) were lysed in buffer containing 50 mM HEPES pH 7.5, 125 mM KOAc, 5 mM MgOAC_2_, 1% Triton X-100, 10% glycerol, 1 mM DTT, and 1× EDTA-free complete protease inhibitors (Roche). Lysate (8 mg/sample) was incubated with 200 µl anti-Flag magnetic beads (Sigma) at 4°C for 90 min. Beads were washed 3× with lysis buffer, 3× with lysis buffer + 400 mM KOAc, and 3× with elution buffer (50 mM HEPES pH 7.5, 0.1 mM EDTA, 100 mM KCl, 25% glycerol, 1 mM DTT), then eluted in 100 mM elution buffer + 1 mg/ml 3× Flag peptide (Sigma) at 4°C for 30 min.

### Figure preparation

All figures containing cryo-EM density were generated with UCSF Chimera (Pettersen et al., 2004) or UCSF ChimeraX (Pettersen et al., 2021). Density was colored using the Color Zone tool in UCSF ChimeraX (Pettersen et al., 2021) with a 3 Å radius. Maps colored by local resolution were visualized in Chimera using LocalRes in RELION 4.0 (Kimanius et al., 2021) (Figure 3—figure supplement 1 and 3). All figures were compiled in Adobe Illustrator (Adobe).

### Rigor and reproducibility

Biological replicates are defined here as independent measurements of physically distinct samples. Technical replicates are defined as repeated measurements of the same physical sample. eEF1A used for smFRET and cryo-EM experiments was purified from two distinct batches of RRL. For smFRET experiments, the number of traces (N) is indicated in each figure panel and all experiments were performed in biological triplicate and were repeated on different days. Error bars in smFRET experiments represent standard errors obtained by bootstrap analysis (1000 samples) of the pooled data from all experimental repeats. The sample size chosen for bootstrap analysis converged to within the obtainable precision for the experimental setup, i.e. larger samples would have incurred more computation time without yielding additional information. For homoprogargyl glycine metabolic labeling assays and apoptosis assays, IC_50_ values and plotted data points are given as the mean of three independent biological determinations ± standard error.

## Data Availability

MATLAB-based software platform for smFRET analysis SPARTAN (Juette et al., 2016) is freely available at http://scottcblanchardlab.com/software. Cryo-EM 3D maps for all structures are available through the Electron Microscopy Data Bank (EMDB) as follows: ternatin-4-stalled elongating rabbit 80S, EMD-27732; didemnin B-stalled human 80S initiation complex, EMD- 27691; ternatin-4-stalled human 80S initiation complex, EMD-27694.

## Acknowledgements

This work was supported by National Institutes of Health grants GM079238 to S.C.B. We thank D. Terry, R. Kiselev and other members of the Blanchard laboratory for their expertise and efforts to enable the single-molecule investigations performed and for their review of the manuscript during the preparation and completion of this research. We acknowledge support from the Single-Molecule Center at St. Jude Children’s Research Hospital. We thank A. Plante at Weill Cornell Medicine for guidance in data processing and high-performance computing. Some of this work was performed at the Simons Electron Microscopy Center and National Resource for Automated Molecular Microscopy located at the New York Structural Biology Center, supported by grants from the Simons Foundation (349247), NYSTAR, and the NIH National Institute of General Medical Sciences (GM103310). We acknowledge specific support from SEMC members B. Carragher, C. Potter, E. Eng, and D. Bobe for guidance with grid preparation and data collection.

## Author Contributions

M.F.J., A.F., M.R.W., and M.H. performed and analyzed smFRET experiments. J.D.C. performed cellular experiments. A.B. and S.S. performed cryo-EM studies on the rabbit 80S ribosome. E.J.R. and A.F. performed cryo-EM studies on the human 80S ribosome. M.F.J., J.D.C., E.J.R., J.T., and S.C.B. wrote the manuscript. E.J.R. created figure panels and illustrations. J.T. and S.C.B. initiated and supervised the project.

## Supplement

**Figure 1––figure supplement 1.**
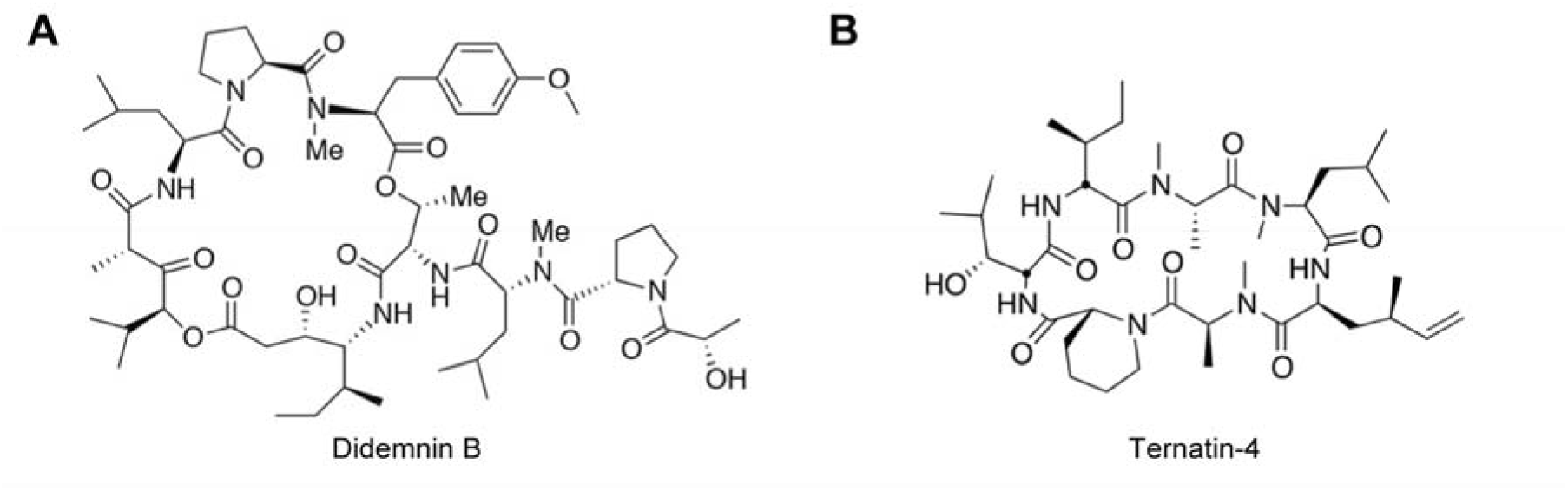
Molecular structures of (**A**) didemnin (didemnin B) and (**B**) ternatin-4.

**Figure 1––figure supplement 2.**
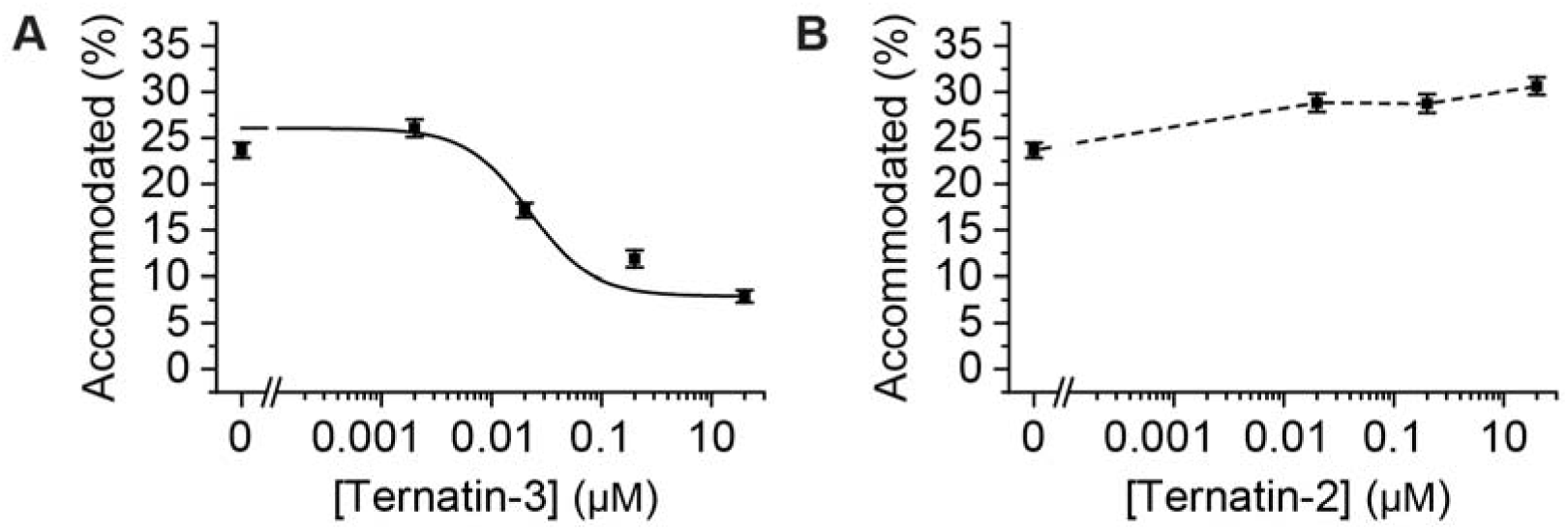
Dose response profiles of the accommodated fraction for (**A**) intermediately active ternatin-3, and (**B**) inactive ternatin-2. Error bars: s.e.m. from 1000 bootstrap samples.

**Figure 1––figure supplement 3.**
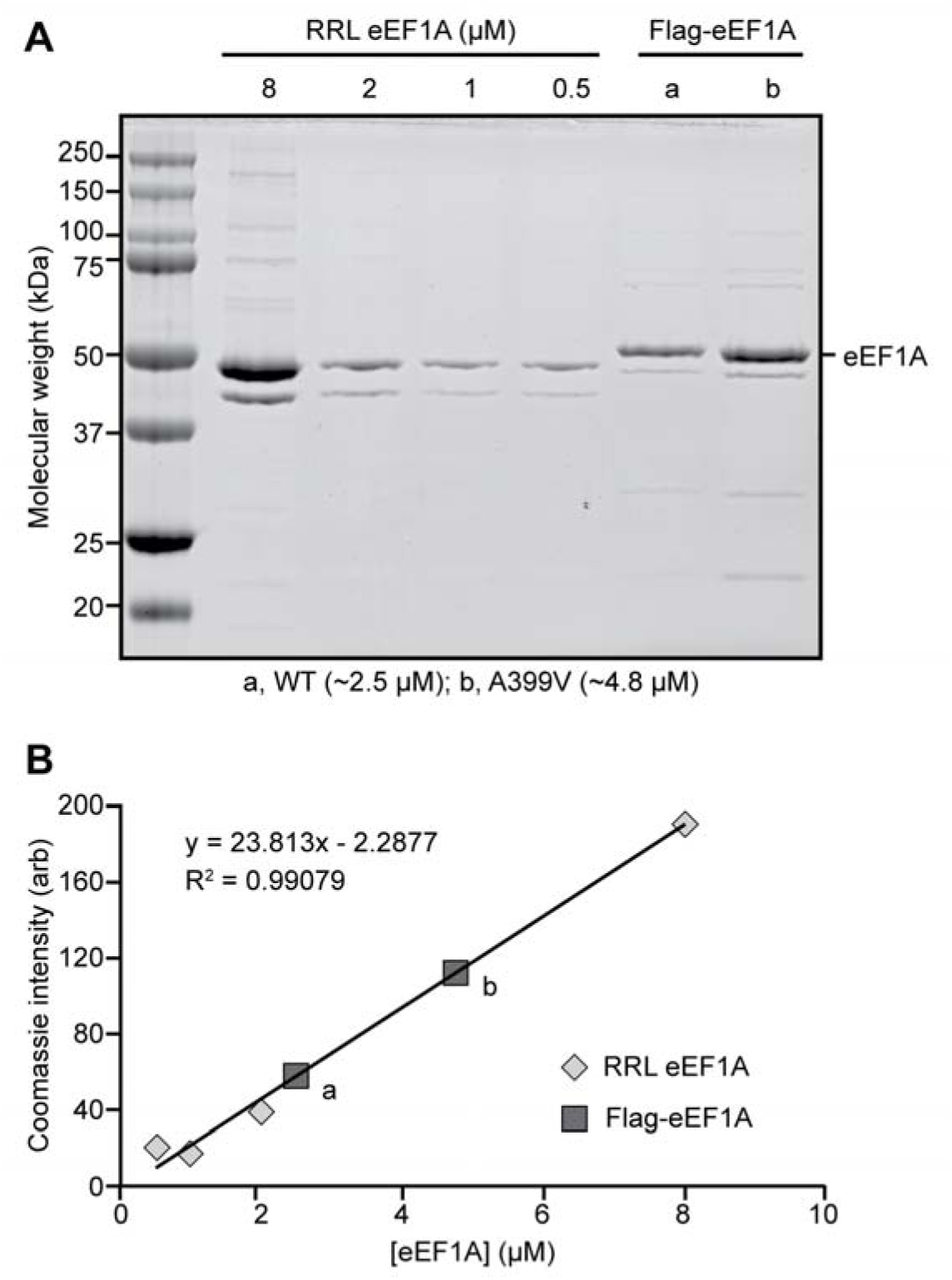
(**A**) Coomassie-stained SDS-PAGE gel loaded with rabbit reticulocyte lysate (RRL) eEF1A (*left*) and wild type (WT; a) or mutant (A399V; b) recombinant Flag-eEF1A (*right*) and (**B**) corresponding concentration calibration plot.

**Figure 2––figure supplement 1.**
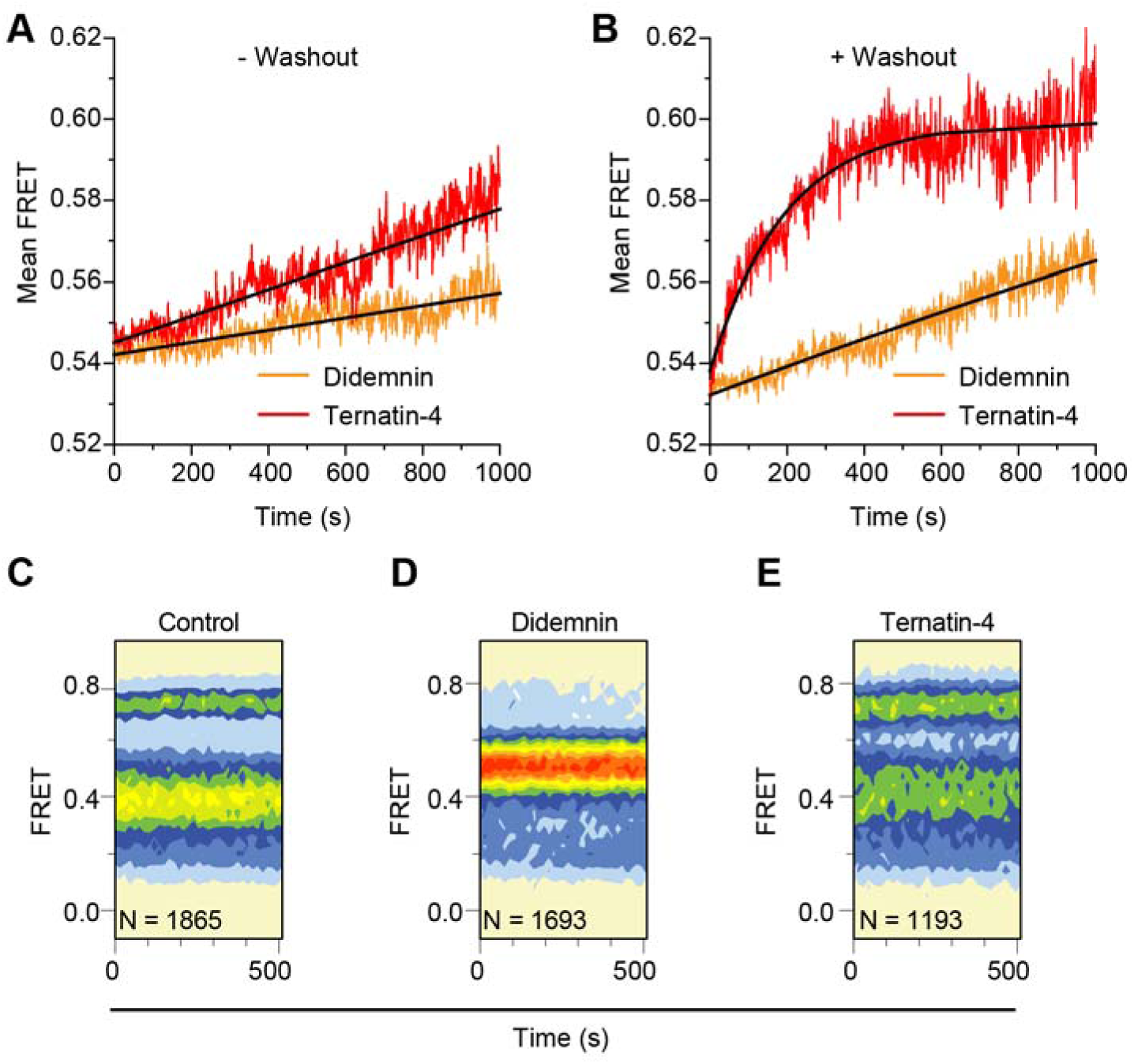
(**A**, **B**) Time course of accommodation in the presence of 350 µM cycloheximide (A) with 20 µM didemnin (orange) or ternatin-4 (red) in solution (black line: linear fits) and (B) after washing out free drug from stalled complexes (black line: exponential fits). (**C**-**E**) Equilibrium population histograms of N traces of (C) drug-free pre-translocation complex, (D) didemnin- or (E) ternatin-4-stalled complexes 5 minutes after washout.

**Figure 3––figure supplement 1.**
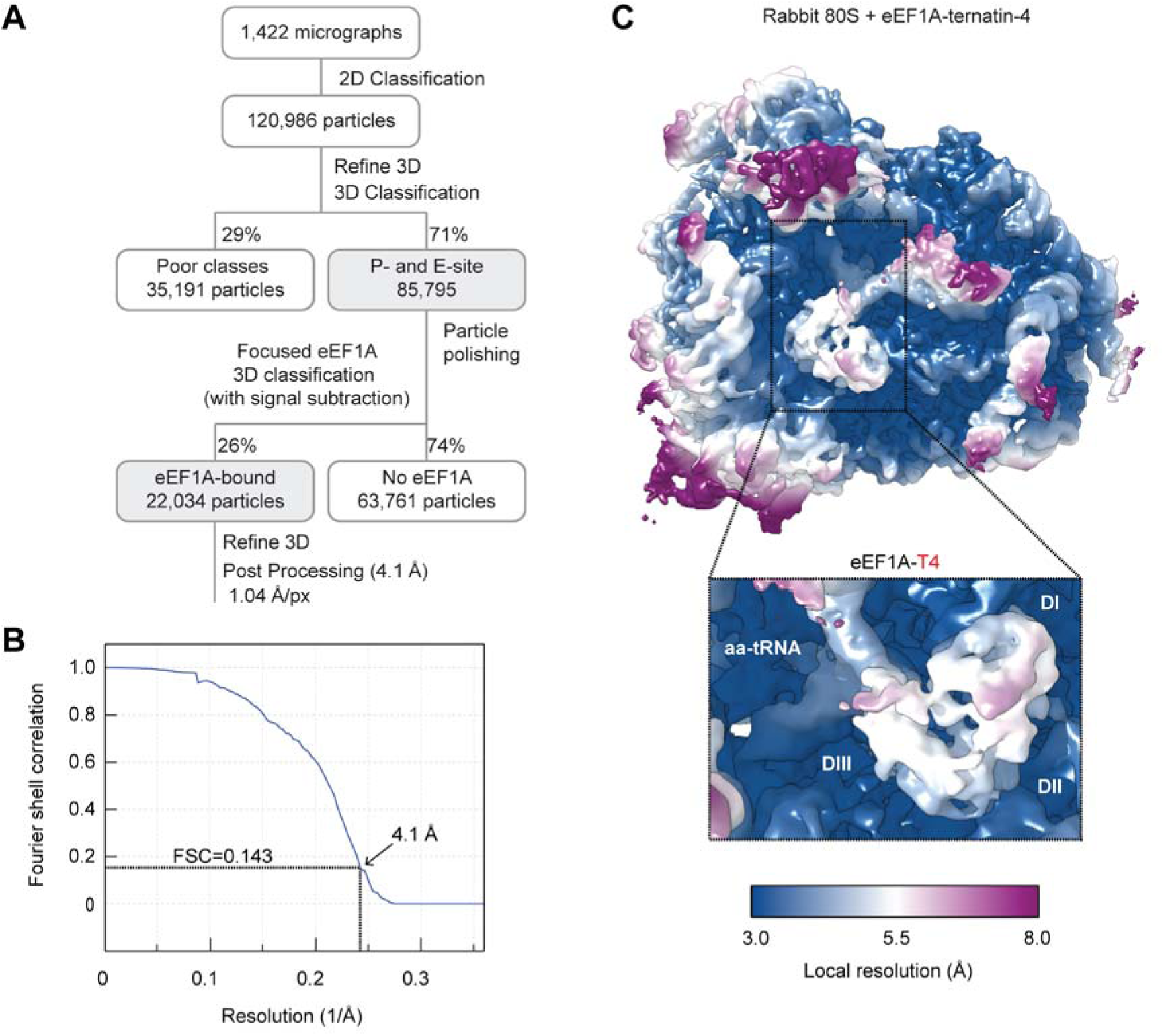
Cryo-EM processing of the ternatin-4 stalled rabbit 80S- eEF1A-aa-tRNA structure. (**A**) Flowchart of cryo-EM image processing. (**B**) Fourier Shell Correlation (FSC) curve for the final reconstruction. Based on the FSC = 0.143 criterion the map reaches a nominal resolution of 4.1 Å. (**C**) Local resolution of refined map showing an overview and eEF1A (*inlay*). The local resolution for the eEF1A(GDP)-aa-tRNA ternary complex is lower than the surrounding ribosome due to flexibility.

**Figure 3––figure supplement 2.**
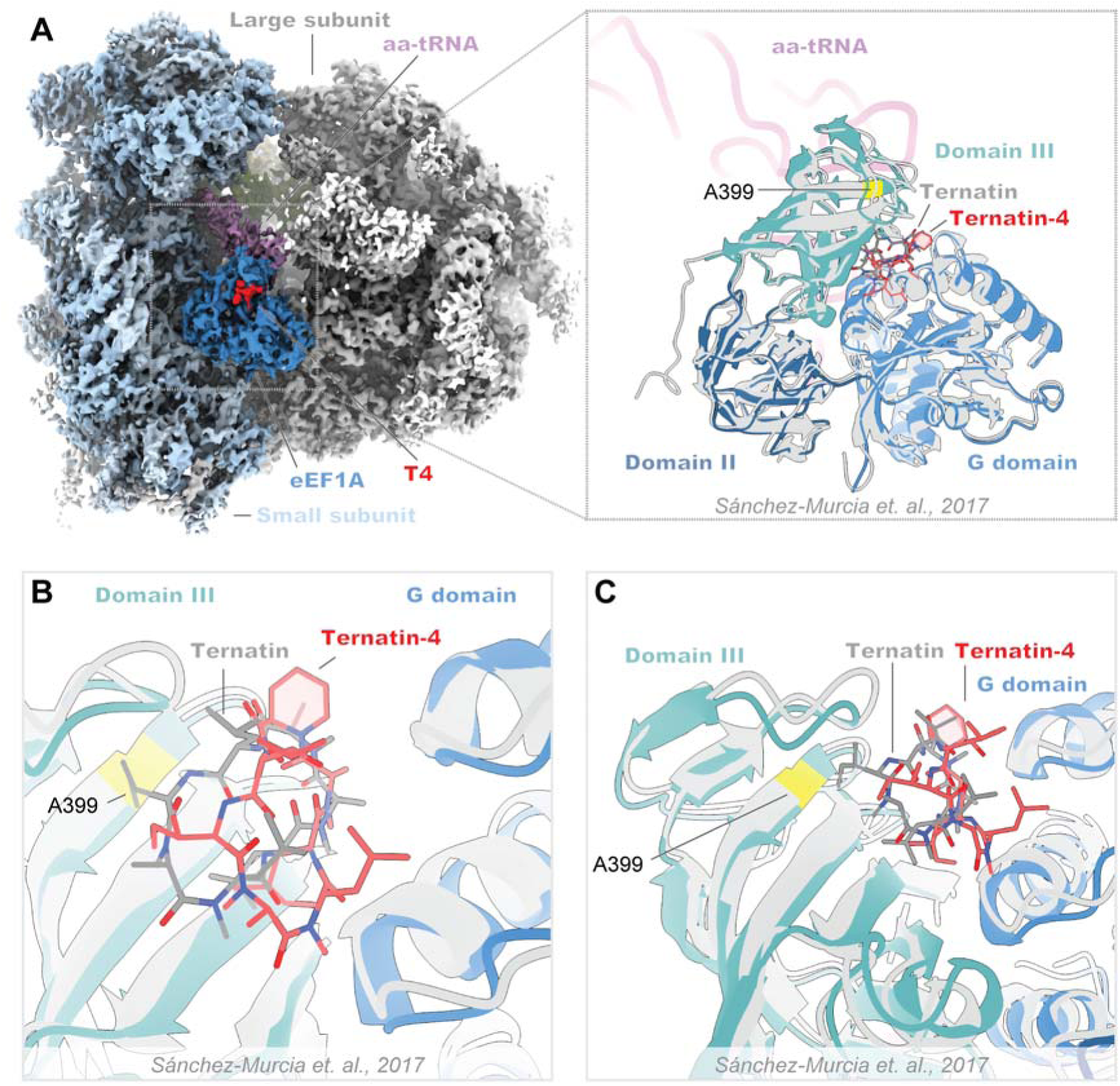
Structural comparison of the ternatin-4-stalled rabbit elongation complex with a published prediction of ternatin binding. (**A**) Overview of the cryo-EM density maps of the ternatin-4-stalled rabbit elongation complex (*left*) comprising the large (LSU; gray) and small (SSU; light blue) ribosomal subunits, peptidyl-tRNA (P site; green), aminoacyl-tRNA in the pre-accommodated A/T state (aa-tRNA; purple), eEF1A (blue), and ternatin-4 (T4; red). (Inlay) Molecular model of eEF1A and aa-tRNA (purple) from PDB-ID: 5LZS (Sánchez-Murcia et al., 2017), colored by domain was rigid-body fit into the cryo-EM map and aligned to a published molecular dynamics model of ternatin docked to eEF1A (gray) (Sánchez-Murcia et al., 2017). (**B**, **C**) Zoom-in of the overlay from panel (A; inlay) of the ternatin- 4 binding site at the interface between the G domain and domain III of eEF1A, colored as in (A; inlay). Residue A399 (yellow), which confers resistance to ternatin and didemnin when mutated to valine, is adjacent to the density for T4.

**Figure 3––figure supplement 3.**
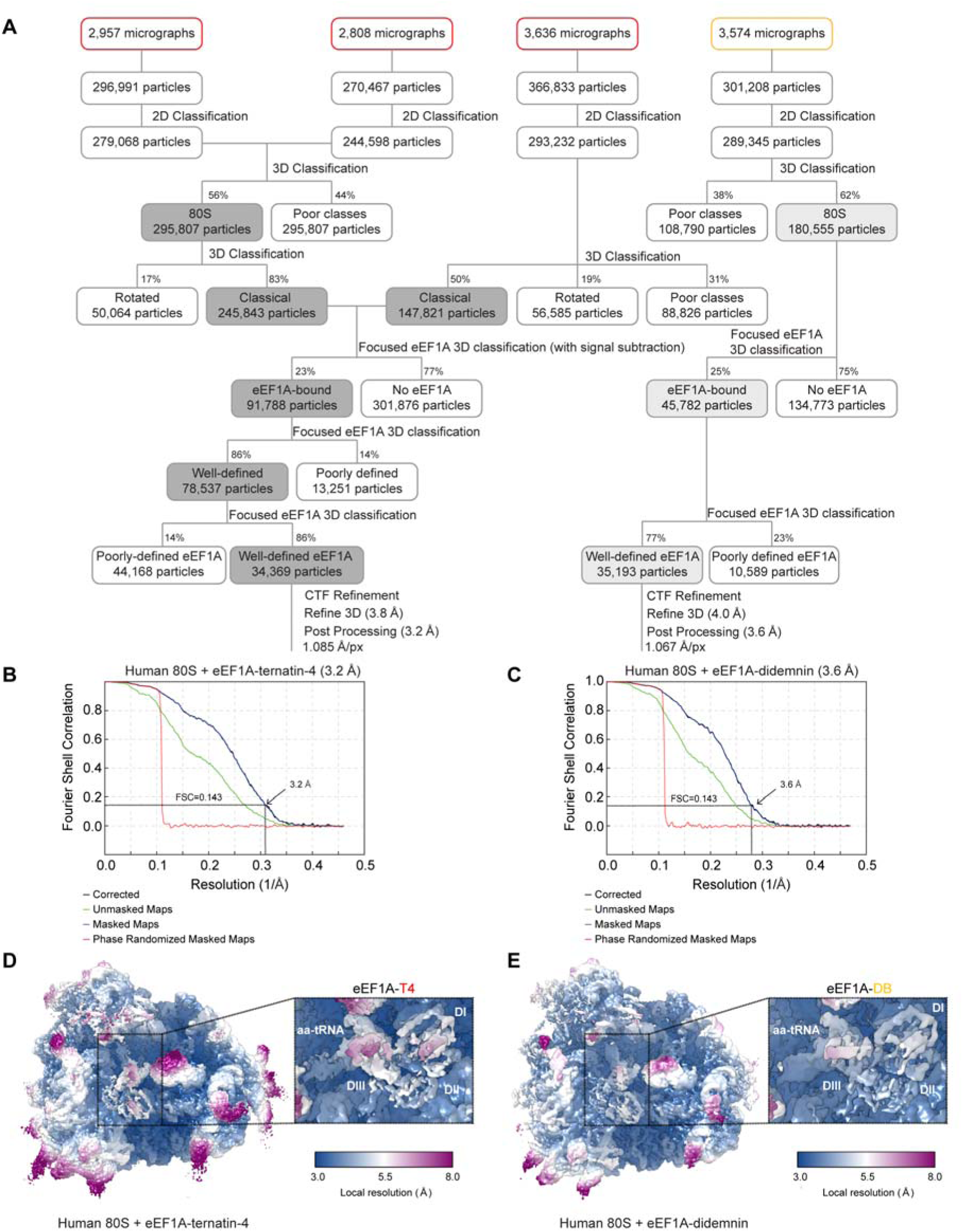
Cryo-EM processing of the didemnin and ternatin-4 stalled human 80S-eEF1A-aa-tRNA structures. (**A**) Flowchart of cryo-EM image processing for final map generation for ternatin-4 (red; *left*) and didemnin (orange; *right*). (**B**, **C**) Fourier Shell Correlation (FSC) curve for the final reconstructions for (B) ternatin-4 and (C) didemnin. (**D**, **E**) Local resolution of the refined maps for (D) ternatin-4 (T4) and (E) didemnin (DB) viewing overview and eEF1A (*inlay*). The local resolution for the eEF1A-aa-tRNA ternary complex is lower than the surrounding ribosome due to flexibility.

**Figure 3––figure supplement 4.**
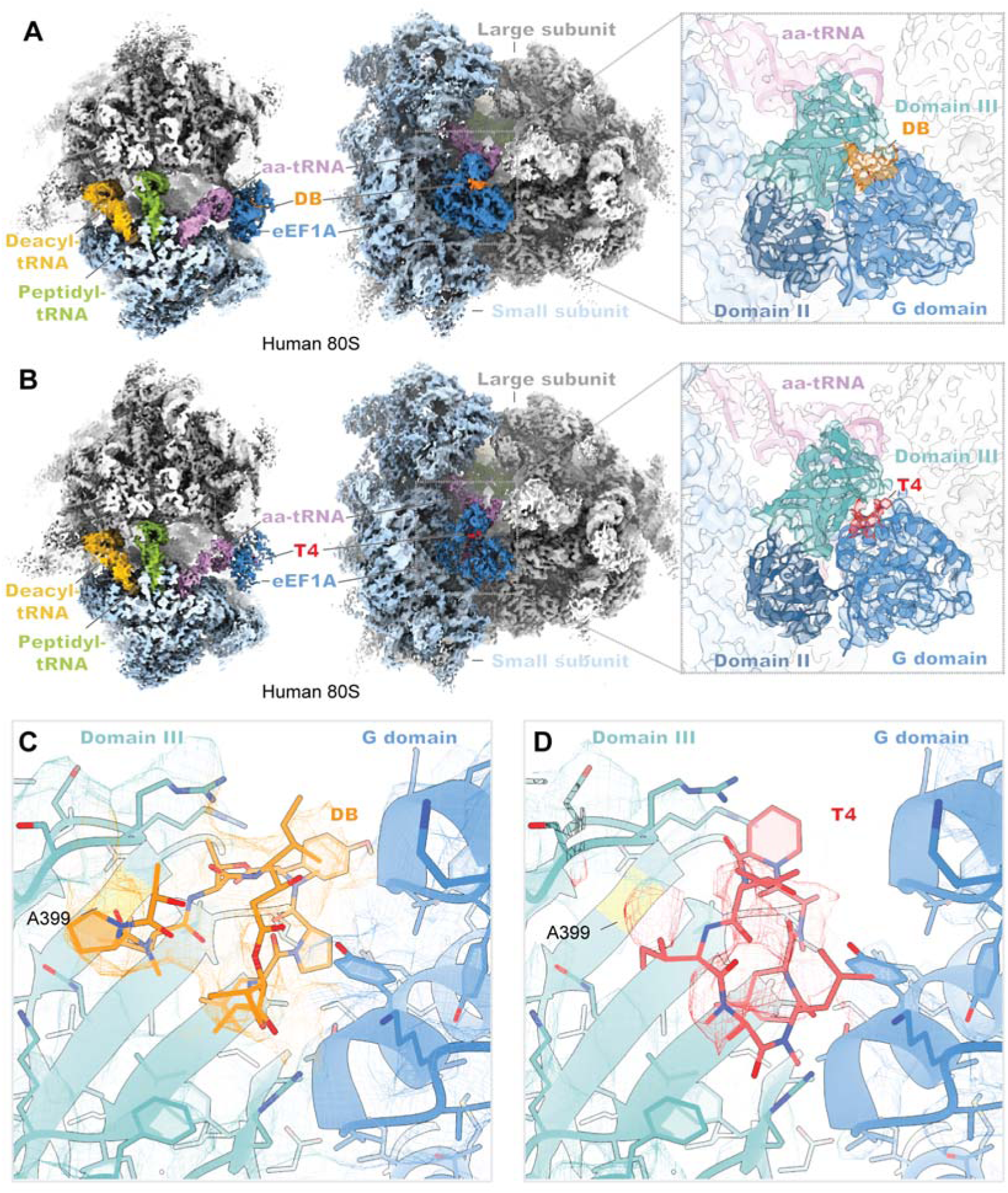
Cryo-EM structures of didemnin and ternatin-4 stalled human 80S -eEF1A(aa-tRNA) complexes. (**A**, **B**) Overview of the cryo-EM density maps of the (A) didemnin- and (B) ternatin-4-stalled human initiation complexes viewed from the small subunit (SSU) head domain (*left*) and into the GTPase activating center (GAC) from the leading edge (*middle*) comprising the large subunit (LSU; gray) and SSU (light blue), peptidyl-tRNA (P site; green) and deacyl-tRNA (E site; gold), aminoacyl-tRNA in the pre-accommodated A/T state (aa-tRNA; purple), eEF1A (blue), and didemnin (DB; orange) or ternatin-4 (T4; red). Inlays (*right*) show cryo-EM density of eEF1A ternary complex on the ribosome highlighting density for (A) DB or (B) T4, with eEF1A colored by domain. Molecular model of eEF1A and aa-tRNA from PDB-ID: 5LZS (Shao et al., 2016) was rigid-body fit into both cryo-EM maps. (**C, D**) Zoom-in of unaccounted cryo-EM density at the interface between the G domain and domain III of eEF1A has been assigned to (C) DB and (D) T4, colored as in (A, B; *left*). Residue A399 (yellow), which confers resistance to ternatin and DB when mutated to valine, is adjacent to the overlapping drug binding pocket. All cryo-EM density is contoured at 2.5 σ.

**Figure 4––figure supplement 1.**
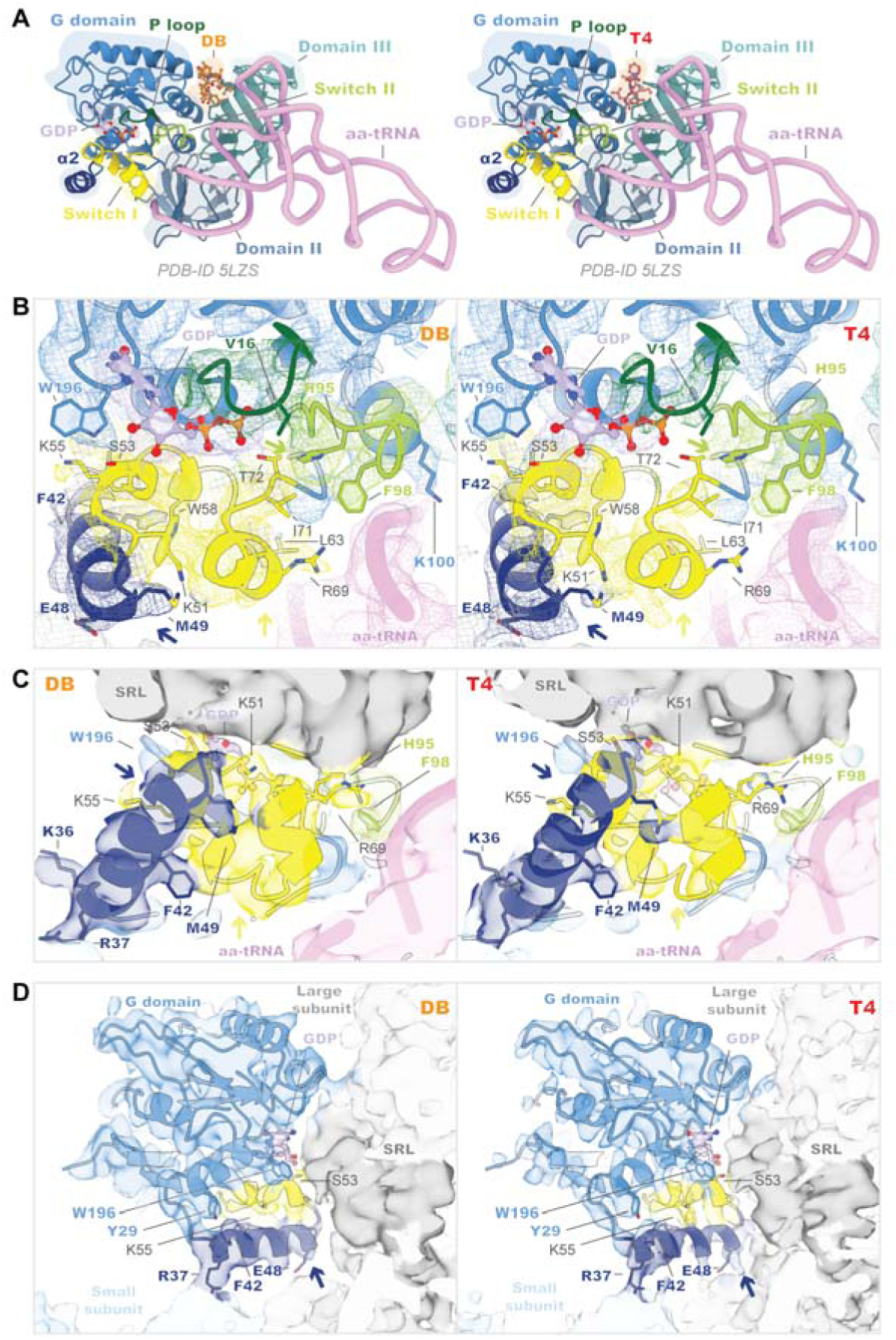
The G domain of eEF1A is more dynamic when bound to ternatin-4 than to didemnin on the human 80S ribosome. (**A**) Overview of the domain architecture of eEF1A ternary complex from PDB-ID: 5LZS (Shao et al., 2016) bound to didemnin (DB; orange; *left*) or ternatin-4 (T4; red; *right*) as viewed from the leading edge of the human 80S ribosome and the Sarcin ricin loop (SRL). G-domain (blue) elements include switch I (yellow), switch II (lime), the P loop (dark green), helix α2 (dark blue), and a bound GDP (light purple) in the nucleotide binding pocket. pre-accommodated A/T state (aa-tRNA; pink), eEF1A (blue), and ternatin-4 (T4; red). (**B**-**D**) Molecular models from PDB-ID: 5LZS (Shao et al., 2016) and cryo-EM density shown in (B) mesh and (C, D) surface representation of the eEF1A G domain when stalled with didemnin (DB; orange; *left*) or T4 (*right*) on the elongating rabbit 80S ribosome, colored as in (A). Colored arrows indicate regions of weakened cryo-EM density in the T4-stalled eEF1A G domain in the C terminus of helix α2 (dark blue), the C terminus of switch I (yellow) and the catalytic His95 of switch II (lime). Panels highlight (B) the nucleotide binding pocket of eEF1A and switch loop architecture, (C) the interface between eEF1A and the SRL (dark gray), and (D) the junction between the SRL, eEF1A helix α2, and small subunit (SSU; light blue) rRNA helix 14. All cryo-EM density is contoured at 2 σ.

**Figure 5––figure supplement 1.**
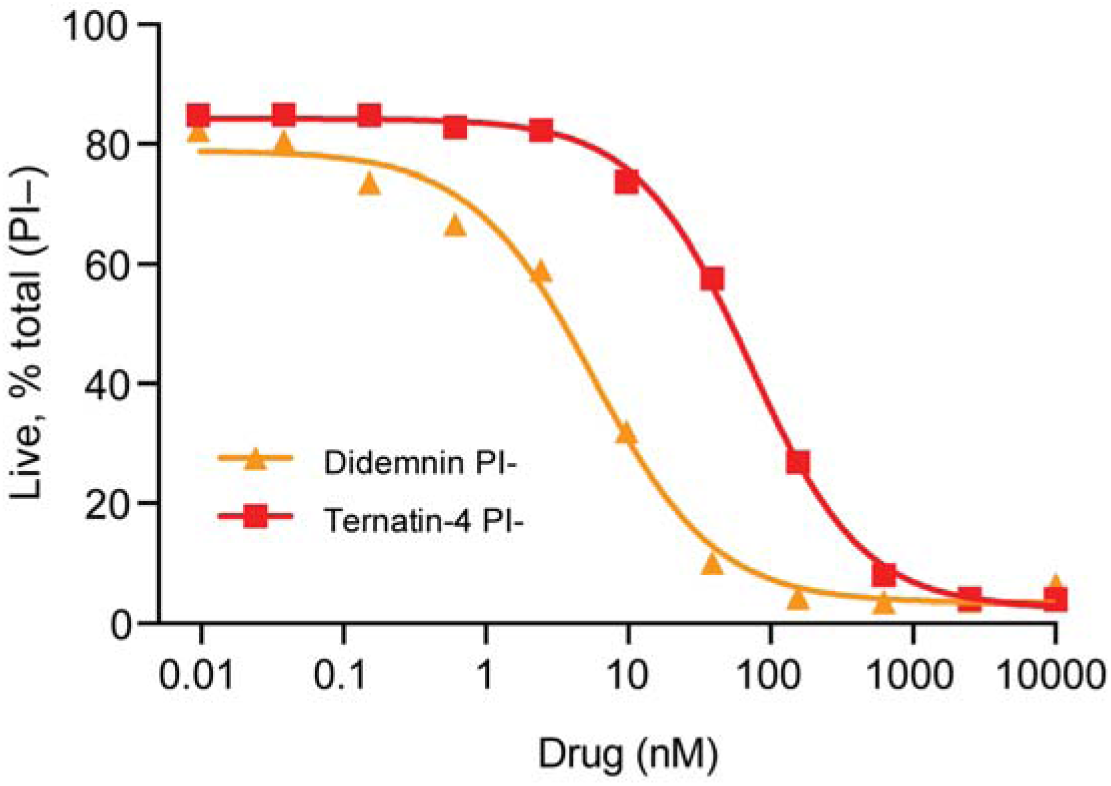
Dose response curve for Jurkat cells treated for 24 hours with the indicated compound. Cells were stained with propidium iodide (PI-) and analyzed for cell death by flow cytometry.

**Table S1.** Apparent transition rates k_i◊j_ between FRET states (low FRET: index 1, intermediate FRET: index 2, high FRET: index 3) and overall decay rates k_i_ for each state observed in aa-tRNA selection experiments prior to the first dwell in high FRET (AC) for ≥ 150 ms with standard errors from 1000 bootstrap samples.

| **Control** |  |  |
| --- | --- | --- |
| $k_1 = (1.90 \pm 0.13) \text{ s}^{-1}$ | $k_{1 \rightarrow 2} = (1.89 \pm 0.13) \text{ s}^{-1}$ | $k_{1 \rightarrow 3} = (0.01 \pm 0.005) \text{ s}^{-1}$ |
| $k_{2 \rightarrow 1} = (0.99 \pm 0.01) \text{ s}^{-1}$ | $k_2 = (1.73 \pm 0.09) \text{ s}^{-1}$ | $k_{2 \rightarrow 3} = (0.74 \pm 0.06) \text{ s}^{-1}$ |
| $k_{3 \rightarrow 1} = (0.24 \pm 0.09) \text{ s}^{-1}$ | $k_{3 \rightarrow 2} = (15.21 \pm 0.44) \text{ s}^{-1}$ | $k_3 = (15.45 \pm 0.43) \text{ s}^{-1}$ |
| **Didemnin** |  |  |
| $k_1 = (2.827 \pm 0.18) \text{ s}^{-1}$ | $k_{1 \rightarrow 2} = (2.824 \pm 0.18) \text{ s}^{-1}$ | $k_{1 \rightarrow 3} = (0.003 \pm 0.003) \text{ s}^{-1}$ |
| $k_{2 \rightarrow 1} = (0.64 \pm 0.03) \text{ s}^{-1}$ | $k_2 = (0.71 \pm 0.03) \text{ s}^{-1}$ | $k_{2 \rightarrow 3} = (0.07 \pm 0.01) \text{ s}^{-1}$ |
| $k_{3 \rightarrow 1} = (0.22 \pm 0.17) \text{ s}^{-1}$ | $k_{3 \rightarrow 2} = (19.1 \pm 1.1) \text{ s}^{-1}$ | $k_3 = (19.3 \pm 1.1) \text{ s}^{-1}$ |
| **Ternatin-4** |  |  |
| $k_1 = (2.36 \pm 0.15) \text{ s}^{-1}$ | $k_{1 \rightarrow 2} = (2.35 \pm 0.16) \text{ s}^{-1}$ | $k_{1 \rightarrow 3} = (0.01 \pm 0.01) \text{ s}^{-1}$ |
| $k_{2 \rightarrow 1} = (0.71 \pm 0.04) \text{ s}^{-1}$ | $k_2 = (0.85 \pm 0.04) \text{ s}^{-1}$ | $k_{2 \rightarrow 3} = (0.14 \pm 0.02) \text{ s}^{-1}$ |
| $k_{3 \rightarrow 1} = (0.52 \pm 0.42) \text{ s}^{-1}$ | $k_{3 \rightarrow 2} = (17.45 \pm 0.81) \text{ s}^{-1}$ | $k_3 = (17.97 \pm 0.83) \text{ s}^{-1}$ |

**Table S2.**
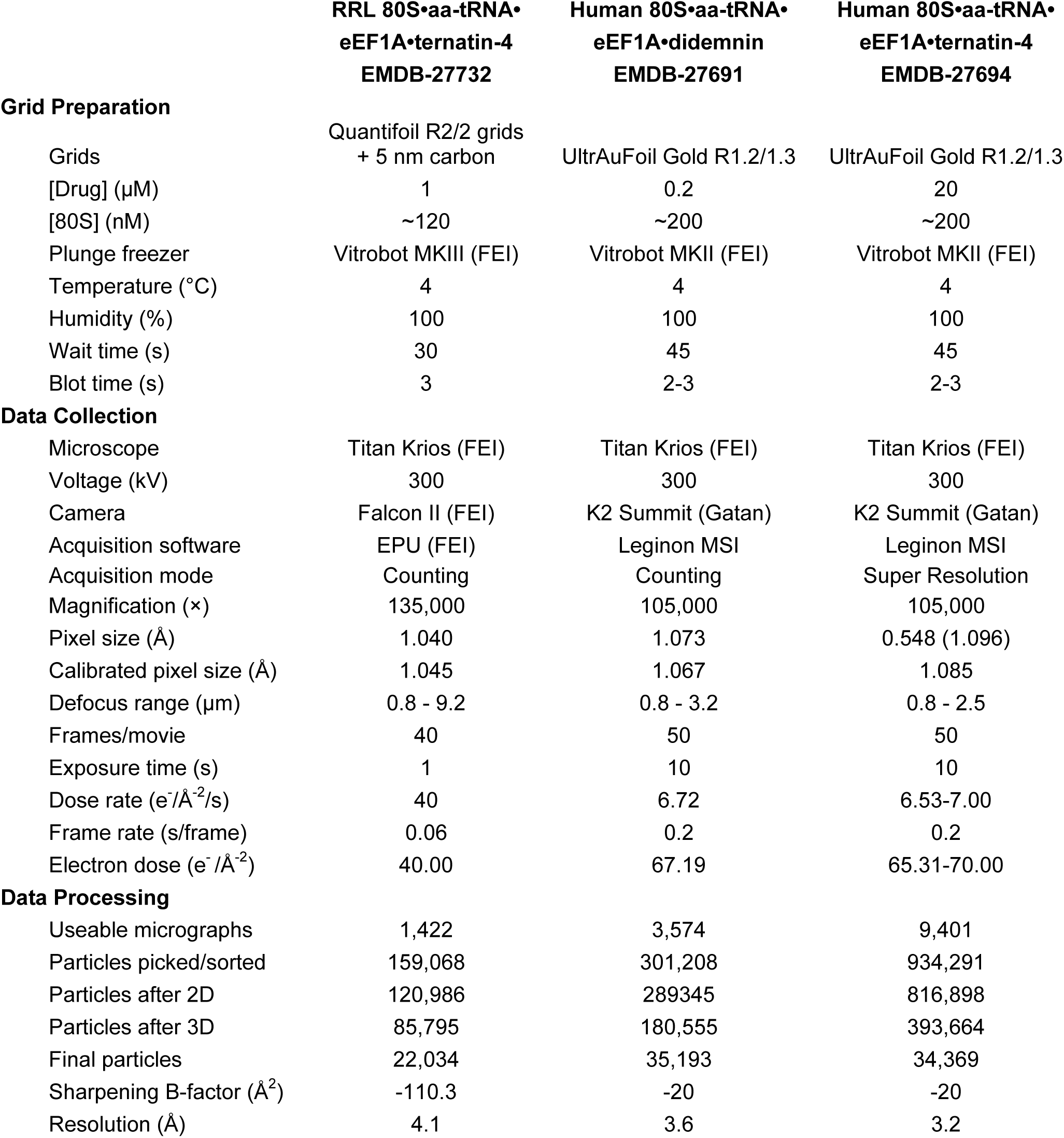
Data collection and processing statistics.

